# Single-cell transcriptional diversity is a hallmark of developmental potential

**DOI:** 10.1101/649848

**Authors:** Gunsagar S. Gulati, Shaheen S. Sikandar, Daniel J. Wesche, Anoop Manjunath, Anjan Bharadwaj, Mark J. Berger, Francisco Ilagan, Angera H. Kuo, Robert W. Hsieh, Shang Cai, Maider Zabala, Ferenc A. Scheeren, Neethan A. Lobo, Dalong Qian, Feiqiao B. Yu, Frederick M. Dirbas, Michael F. Clarke, Aaron M. Newman

**Author notes:** These authors contributed equally (Gunsagar S. Gulati, Shaheen S. Sikandar).

## Abstract

Single-cell RNA-sequencing (scRNA-seq) is a powerful approach for reconstructing cellular differentiation trajectories. However, inferring both the state and direction of differentiation without prior knowledge has remained challenging. Here we describe a simple yet robust determinant of developmental potential—the number of detectably expressed genes per cell— and leverage this measure of transcriptional diversity to develop a new framework for predicting ordered differentiation states from scRNA-seq data. When evaluated on ~150,000 single-cell transcriptomes spanning 53 lineages and five species, our approach, called CytoTRACE, outperformed previous methods and ~19,000 molecular signatures for resolving experimentally-confirmed developmental trajectories. In addition, it enabled unbiased identification of tissue-resident stem cells, including cells with long-term regenerative potential. When used to analyze human breast tumors, we discovered candidate genes associated with less-differentiated luminal progenitor cells and validated *GULP1* as a novel gene involved in tumorigenesis. Our study establishes a key RNA-based correlate of developmental potential and provides a new platform for robust delineation of cellular hierarchies (https://cytotrace.stanford.edu).

## Main Text

In multicellular organisms, tissues are hierarchically organized into distinct cell types and cellular states with intrinsic differences in function and developmental potential^1^. Common methods for studying cellular differentiation hierarchies, such as lineage tracing and functional transplantation assays, have revealed detailed roadmaps of cellular ontogeny at scales ranging from tissues and organs to entire model organisms^2–4^. However, despite the power of these technologies, they cannot be applied to human tissues *in vivo* and generally require prior knowledge of cell type-specific genetic markers^2^. These limitations have made it difficult to study the developmental organization and cell fate decisions of primary human tissues under normal physiological conditions and during disease.

Single-cell RNA-sequencing (scRNA-seq) has recently emerged as a promising approach to study tissue architecture^5,6^ and cellular differentiation trajectories at high resolution in primary tissue specimens^7^. Although a large number of computational methods for predicting lineage trajectories have been described, they generally rely upon (1) *a priori* knowledge of the starting point (and thus, direction) of the inferred biological process^8–14^ and (2) the presence of intermediate cell states to reconstruct the trajectory^15,16^. These requirements, although reasonable in well-established systems and in time-series experiments, can be challenging to satisfy in tissues with poorly understood developmental biology, such as human neoplasms^17^. Moreover, it remains difficult to distinguish quiescent (non-cycling) adult stem cells with longterm regenerative potential from more specialized cells using existing *in silico* approaches. While gene expression-based models can potentially overcome these limitations (e.g., transcriptional entropy^18–20^, pluripotency-associated gene sets^21^ and machine learning strategies^22^), their relative utility across diverse developmental systems and single-cell sequencing technologies is still unclear.

Here, we profiled nearly 19,000 features of single-cell gene expression data to discover factors that accurately predict cellular differentiation states independently of tissue type, species, and platform. Among the top-performing features, we identified a simple yet surprisingly effective determinant of developmental potential—the number of detectably expressed genes per cell. By leveraging this measure of transcriptional diversity, which was noisy at the single-cell level, we developed a new unsupervised framework for determining ordered differentiation states from single-cell transcriptomes, called CytoTRACE (Cellular (<u>Cyto</u>) <u>T</u>rajectory <u>R</u>econstruction <u>A</u>nalysis using gene <u>C</u>ounts and <u>E</u>xpression). We show that our approach (1) substantially outperforms leading computational methods and 18,706 molecular signatures for predicting differentiation states in 53 experimentally-confirmed developmental trajectories, (2) reveals cellular hierarchies in whole tissues and whole organisms, and (3) identifies key genes associated with stemness and differentiation in both healthy tissues and human cancer. Our results suggest that CytoTRACE can complement existing lineage trajectory tools and aid the identification of immature cells in diverse multicellular systems.

## Results

### RNA-based correlates of single-cell differentiation states

We sought to identify robust, RNA-based determinants of developmental potential without the need for *a priori* knowledge of developmental direction or intermediate cell states marking cell fate transitions. Toward this end, we evaluated ~19,000 potential correlates of cell potency in scRNA-seq data, including all available gene sets in the Molecular Signatures Database (*n* = 17,810)^23^, 896 gene sets covering transcription factor binding sites from ENCODE^24^ and ChEA^25^, an mRNA-expression-derived stemness index (mRNAsi)^22^, and three computational techniques that infer stemness as a measure of transcriptional entropy (StemID, SCENT, SLICE^18–20^). We also explored the utility of ‘gene counts’, or the number of detectably expressed genes per cell, which has been anecdotally observed to correlate with differentiation status^26–28^, but not yet comprehensively evaluated (**Methods**).

To assess these features, we compiled a training cohort consisting of nine gold standard scRNA-seq datasets with experimentally-confirmed differentiation trajectories. These datasets were selected to prioritize commonly used benchmarking datasets from prior studies^12,18–20,26,29^ and to ensure a broad sampling of unique developmental states from the mammalian zygote to terminally differentiated cells^26,30^. Overall, the training cohort encompassed 3,174 single cells spanning 49 phenotypes, six tissue types, and three scRNA-seq platforms (**Fig. 1A; Methods**). To determine performance, the mean value of each feature for all previously annotated cellular phenotypes was calculated and correlated against ground-truth differentiation states. The resulting coefficients (Spearman) were then averaged across the nine training datasets to yield a final score and rank for every feature (**Fig. 1B; Methods**).

**Figure 1.**
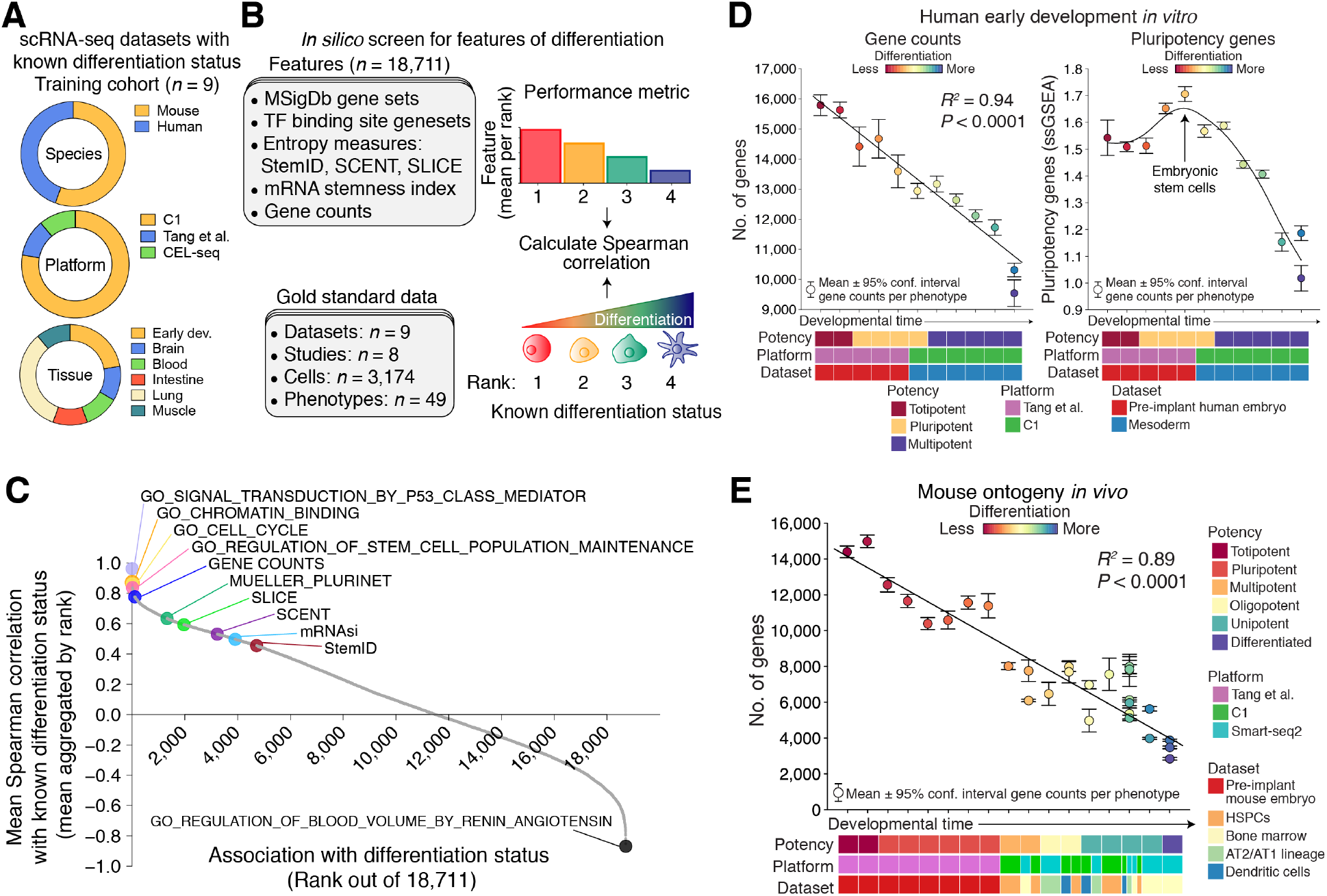
RNA-based determinants of developmental potential. (**A-C**) *In silico* screen for correlates of cellular differentiation status in scRNA-seq data. (**A**) Composition of the training cohort. (**B**) *Left:* Summary of evaluated features (top) and gold standard scRNA-seq datasets from the training cohort (bottom). In total, 17,810 gene sets from MSigDb, 896 transcription factor (TF) binding sites from ENCODE and ChEA, three measures of transcriptional entropy (StemID, SCENT, and SLICE), a machine learning model (mRNAsi), and gene counts (number of detectably expressed genes per cell) were assessed. *Right:* Depiction of the scoring scheme. Each phenotype was assigned a rank based on its known differentiation status (less differentiated = lower rank) and the values of each feature were mean-aggregated by rank for each dataset (higher value = lower rank). Performance was calculated as the mean Spearman correlation between known and predicted ranks across all nine training datasets. (**C**) Performance of all features for predicting differentiation states in the training cohort, ordered by mean Spearman correlations. (**D**) The developmental ordering of 12 human cell phenotypes during early embryogenesis, shown as a function of single-cell gene counts (*left*) and the expression of pluripotency genes^21^ (*right*). Points and error bars denote means and 95% confidence intervals, respectively. Phenotypes are ranked according to their known developmental potential relative to other cell types (phenotype labels are provided in **Methods**). The coefficient of determination (*R^2^*) is shown for gene counts (*left*) and a smooth spline is shown for pluripotency gene enrichment scores (*right*). ssGSEA, single sample gene set enrichment analysis. (**E**) Same as in **D** (left panel) but showing the ontological ordering of 30 mouse cell phenotypes across 17 developmental stages versus single-cell gene counts. For further details and information on datasets, see **Methods**.

This systematic screen revealed many known and unexpected correlates of differentiation status (**Fig. 1C; Fig. S1A**). However, one feature in particular showed surprisingly strong performance – the number of detectably expressed genes per cell (‘gene counts’). Appearing in the top 1% of the ranked list (104 out of 18,711), this data-driven feature compared favorably to well-established stemness programs, including cell cycle and pluripotency signatures^21,22^, yet also showed evidence of unique biology and broader applicability. For example, regardless of whether we examined cycling cells, non-cycling cells, or the earliest stages of human embryogenesis prior to the upregulation of pluripotency factors, gene counts generally decreased with successive stages of differentiation (**Fig. 1D, left; Fig. S2**). Pluripotency genes, by contrast, showed an arc-like pattern during human development, characterized by progressively increasing expression until the emergence of embryonic stem cells, followed by decreasing expression (**Fig. 1D, right**).

These findings suggested that gene counts might extend beyond isolated experimental systems to recapitulate the full spectrum of cellular ontogeny. To formally test this possibility, we compiled, remapped, and normalized a set of mouse lineage trajectories profiled *in vivo* by five plate-seq experiments encompassing 5,059 cells and 30 phenotypes that together span all major potency levels^31^ (**Methods**). Indeed, when averaged by known phenotypes and assessed across independent studies, the relationship between gene counts and differentiation was robustly maintained (*R^2^* = 0.89, *P* < 0.0001; **Fig. 1E; Methods**).

Given these striking results, we performed a series of experiments to better understand the biological basis of gene counts and the factors that influence its measurement.

### Robustness and biological basis of gene counts

We started by characterizing the robustness of gene counts to variation in two key technical parameters: (1) sparsity in single-cell gene expression data and (2) the number of sequenced reads per cell. To investigate the former, we compared gene counts derived from single-cell transcriptomes with gene counts derived from bulk RNA-seq profiles^32^, pooled single-cell transcriptomes, and single-cell transcriptomes following missing value imputation^33^. Regardless of the approach, we observed significantly reduced performance for predicting differentiation states when attempting to overcome sparsity (**Fig. S3A-D**). This suggests that sparsity in scRNA-seq data is driven by real biological heterogeneity in addition to technical noise. Such heterogeneity, while informative for gene counts as a measure of developmental potential, is lost or severely degraded at the population level (**Fig. S3B, E**).

We next examined the relationship between gene counts and the number of reads per cell. Reanalysis of seven scRNA-seq experiments profiled by plate-based protocols showed that even when down-sampling to 10,000 reads per cell, the predictive performance of gene counts was largely maintained (**Fig. S4A-C**). Moreover, the mean number of detectably expressed genes per dataset was linearly related to the logarithm of the mean number of reads (mean *R^2^* = 0.99; **Fig. S4D**). As a result, for most datasets, variation in gene counts due to fluctuations in the number of reads was minimal. Furthermore, predictive performance was only modestly impacted when varying the expression threshold for calculating the number of expressed genes per cell and was unaffected by the removal of potential doublets (**Fig. S5A, B**).

To investigate potential biological correlates of gene counts, we next compared it with the number of detectable mRNA molecules per cell, as measured by unique molecular identifiers (UMIs), external spike-in standards (ERCCs), and Census, a statistical approach to infer the number of mRNA transcripts that are available for capture following cell lysis^34^. By analyzing UMI (*n* = 14 datasets) and ERCC (*n* = 7 datasets) data from previously published droplet-based and plate-based experiments, respectively, we found that a large proportion of the variance in gene counts could be attributed to single-cell mRNA content alone (UMI: mean *R^2^* = 0.84; ERCC: mean *R^2^* = 0.29; **Fig. S6A, B**). This relationship was further confirmed in 17 non-UMI datasets that lack external standards using Census^34^, which produced estimates that were nearly perfectly correlated with the number of unique detectable genes (i.e., canonical transcripts) per cell (mean *R^2^* ≈ 1) (**Fig. S6C; Methods**). We also measured the linear association between gene counts and the number of unique protein-coding splice isoforms per cell. As expected, across 10 plate-seq datasets, gene counts and mRNA diversity were tightly interrelated (mean *R^2^* = 0.98; **Fig. 2A; Fig. S6D**).

**Figure 2.**
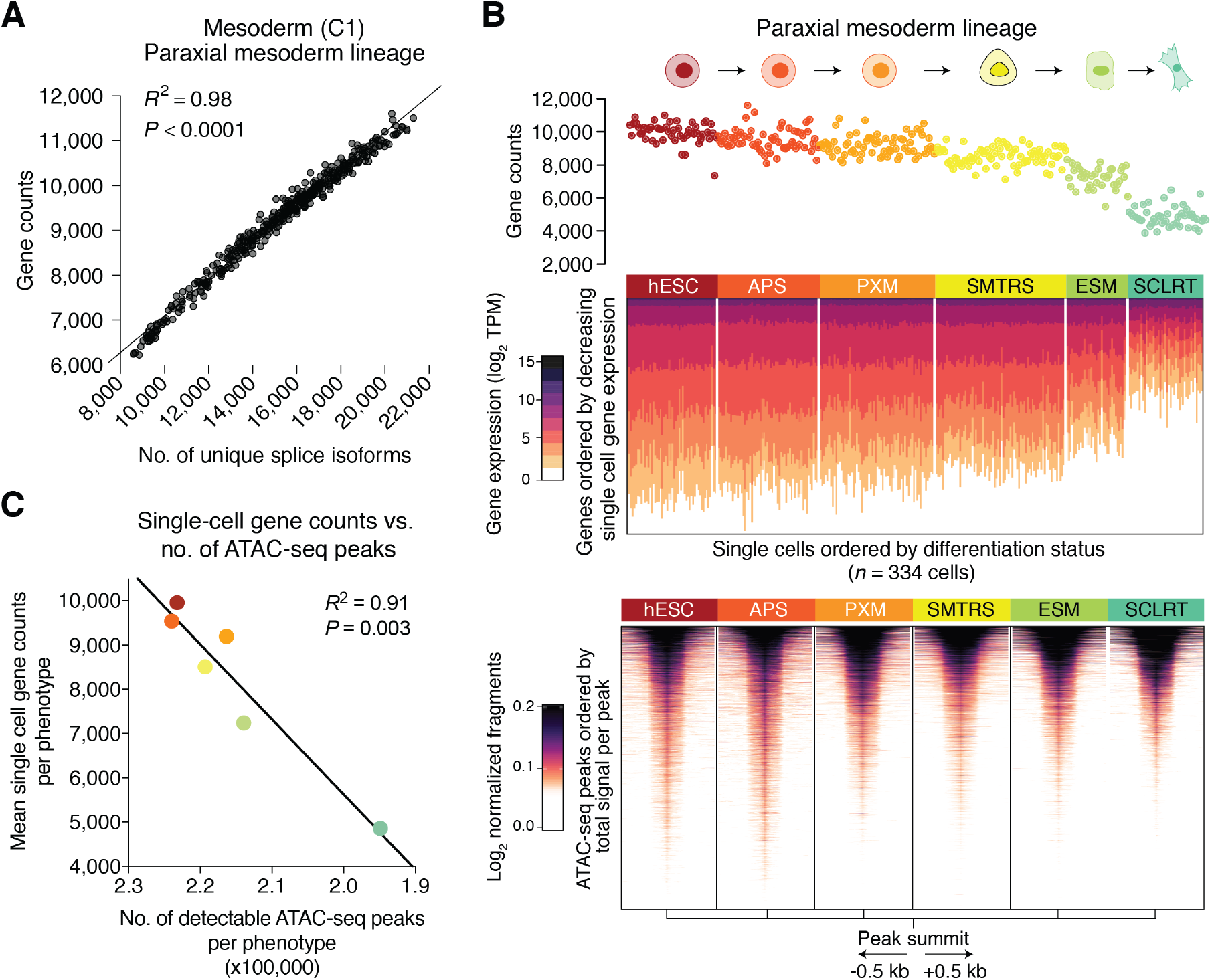
Association between gene counts, RNA diversity, and chromatin accessibility. (**A-C**) Analysis of the association between single-cell gene counts, transcriptional diversity, and chromatin accessibility in cells from an *in vitro* differentiation series of purified phenotypes from the human paraxial mesoderm lineage (**Methods**; also see **Figure S7**). (**A**) Scatterplot showing the association of gene counts with the number of unique protein-coding splice isoforms detected per cell. (**B**) *Top:* Scatterplot showing the association of gene counts (*y*-axis) with paraxial mesoderm differentiation at the single-cell level (*x*-axis). Each point represents a cell colored by known phenotype (below). *Center:* Heat map depicting each single-cell transcriptome in the above scatterplot, but ordered from top-to-bottom by decreasing gene expression (log_2_ TPM). Cells are ordered left to right by increasing differentiation status. *Bottom:* Heat map showing chromatin accessibility profiles (bulk ATAC-seq) for the same cell phenotypes as above. Peaks are centered by their summit, defined as the base with maximum coverage, shown within a window of 1 kb (±0.5 kb), and ordered top to bottom within each phenotype by decreasing total signal per peak. (**C**) Scatterplot showing the concordance between the average number of single-cell gene counts per phenotype and the number of ATAC-seq peaks per phenotype. Points indicate cell types and are colored as in **B**. Linear relationships in **A and C** were evaluated by linear regression (*R^2^*) and corresponding *P* values were determined by a *t*-test. hESC, human embryonic stem cell; APS, anterior primitive streak; PXM, paraxial mesoderm; SMTRS, somitomeres; ESM, early somites; SCLRT, sclerotome.

Since the transcriptional output of a cell is associated with its genome-wide chromatin profile, we hypothesized that single-cell gene counts might ultimately be a surrogate for global chromatin accessibility, which has been previously shown to decrease with differentiation^35–38^. To test this, we compared single-cell gene counts derived from scRNA-seq data with paired bulk ATAC-seq (assay for transposase-accessible chromatin sequencing) profiles obtained from a recent study of *in vitro* mesodermal differentiation from human embryonic stem cells (hESCs)^32^ (**Fig. 2B; Fig. S7A**). In support of our hypothesis, genome-wide chromatin accessibility was observed to progressively decrease with differentiation of hESCs into paraxial mesoderm and lateral mesoderm lineages (**Fig. 2B; Fig. S7A**). Moreover, when segregated by developmental lineage, we observed strong concordance between the number of accessible peaks and the mean number of detectably expressed genes per phenotype (**Fig. 2C; Fig. S7B, C**).

### Development of CytoTRACE

Although gene counts generally showed robust performance when averaged by known phenotypes, in some datasets, such as the *in vitro* differentiation of hESCs into the gastrulation layers^39^, it exhibited considerable intra-phenotypic variation (**Fig. 3A, left**). In fact, when evaluated at a single-cell level, 412 predefined gene sets from our *in silico* screen outperformed gene counts (**Fig. S1B**). Since scRNA-seq was designed to capture single-cell gene expression, we reasoned that genes whose expression patterns correlate with gene counts might better capture differentiation states. Remarkably, by simply taking the geometric mean of genes that were most correlated with gene counts in each dataset (**Fig. S8A-C; Methods**), the resulting dataset-specific ‘gene counts signature’ (GCS) became the top-performing measure in the screen, outranking every pre-defined molecular profile and computational tool that we assessed (**Fig. S1B**).

**Figure 3.**
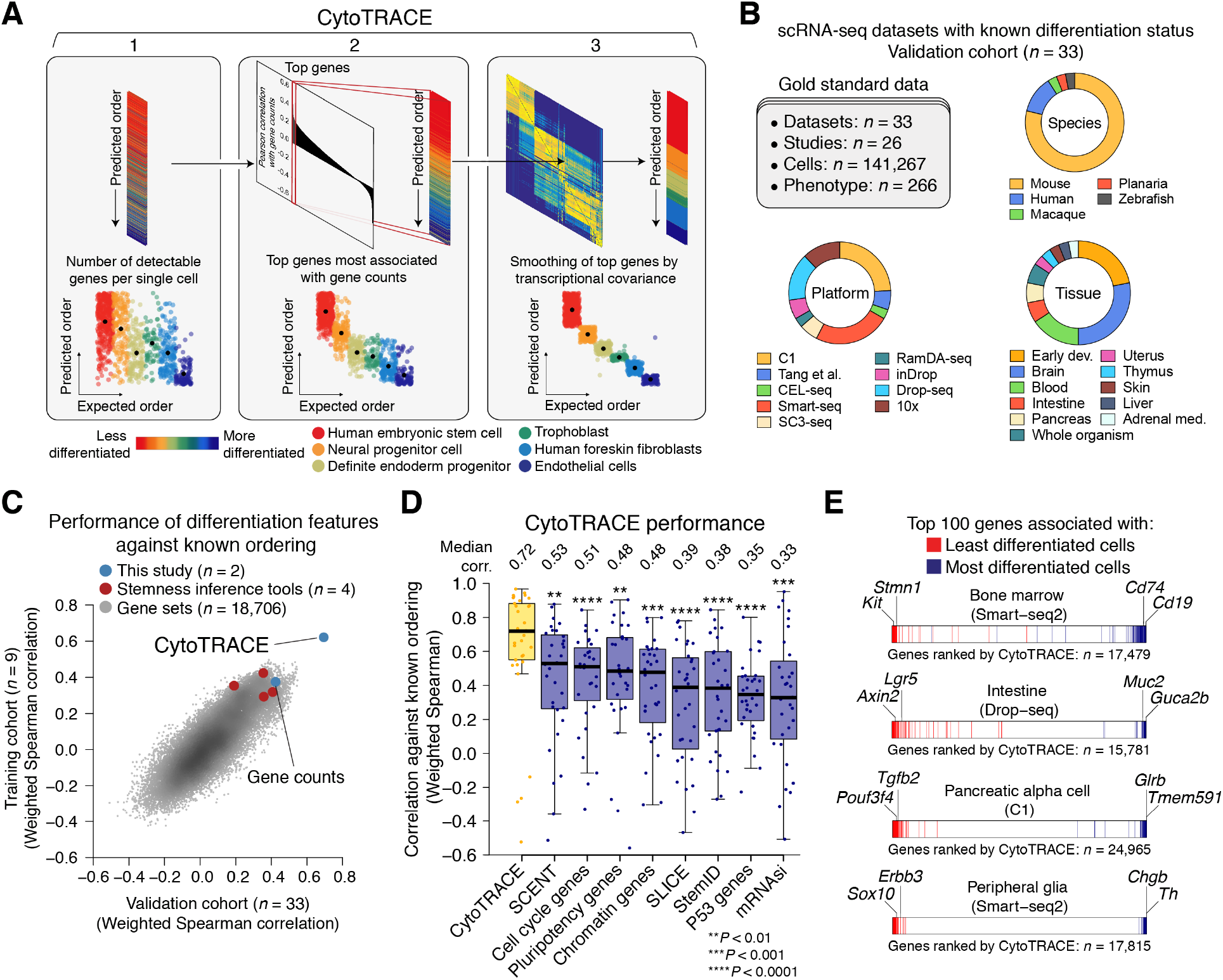
Development and validation of CytoTRACE. (**A**) Schematic overview of the CytoTRACE framework applied to the *in vitro* differentiation of hESCs into the gastrulation layers^39^ (**Methods**). (**B-D**) Validation and benchmarking of CytoTRACE. (**B**) Composition of the validation cohort. In total, 33 scRNA-seq datasets with known differentiation states were analyzed. (**C**) Scatterplot showing predictive performance of 18,706 gene sets, four stemness inference methods^18–20,22^, gene counts, and CytoTRACE, in the training cohort (*y*-axis) and validation cohort (*x*-axis). Results reflect the average single-cell performance per cohort. (**D**) Boxplots showing the single-cell level performance of CytoTRACE against the features and methods from **Fig. 1B** in the validation cohort (*n* = 33 datasets). Each point represents the Spearman correlation, weighted by number of cells per phenotype, between predicted and known differentiation states for a given dataset, calculated as described in **Methods**. Features and computational methods are ordered from left to right by median. Statistical significance was assessed by a one-sided paired Wilcoxon signed-rank test against CytoTRACE. (**E**) Developmental marker gene prioritization without prior knowledge of cellular phenotypes. Plots showing the enrichment of key stemness-associated (red) and differentiation-associated (blue) genes by CytoTRACE in bone marrow (*n* = 4,897 cells), intestine (*n* = 7,216 cells), pancreatic alpha cells; (*n* = 338 cells), and peripheral glia (*n* = 369 cells) (source data are described in **Methods**). Select markers of differentiation are indicated for each dataset. Also see **Figure S12**.

Although GCS is likely to be influenced by multi-lineage priming in some settings^40^, it is derived from all detectably expressed genes per cell in a given dataset. It is therefore inherently robust to drop-out events, agnostic to prior knowledge of developmentally regulated genes, and not solely attributable to a previously defined molecular signature (e.g., pluripotency; **Fig. 1D**). Nevertheless, GCS was still moderately noisy in some datasets (e.g., **Fig. 3A, center; Fig. S8A-C**). We therefore implemented a novel two-step procedure to directly smooth GCS on the basis of transcriptional covariance among single cells (**Fig. 3A, right; Fig. S8A-D; Methods**). The resulting method, which we call CytoTRACE (Cellular (<u>Cyto</u>) <u>T</u>rajectory <u>R</u>econstruction <u>A</u>nalysis using gene <u>C</u>ounts and <u>E</u>xpression), not only significantly outperformed GCS and gene counts (**Fig. S8A**), but also outperformed all evaluated features by a considerable margin (**Fig. S1B**).

### Performance evaluation across tissues, species, and platforms

To validate our findings, we assembled a greatly expanded compendium of 33 additional scRNA-seq datasets. These data were selected to represent diverse experimentally-confirmed developmental lineages and consisted of 141,267 single cells spanning 266 phenotypes, 11 tissue types, five species, nine scRNA-seq platforms (three droplet-based and six plate-based protocols, ranging from an average of ~10,000 to ~1M UMIs or reads per cell, respectively; **Fig. S4A**), and 26 studies (**Fig. 3B; Methods**). As before, performance was determined using Spearman correlations to determine concordance against ground truth data at the single-cell level and by phenotype (**Methods**).

When assessed at the single-cell level, CytoTRACE markedly outperformed existing methods and gene sets (**Fig. 3C, D; Fig. S9A**), and was positively correlated with the direction of differentiation in 88% of datasets (*P* = 7 x 10^−7^, Binomial test). These results were consistent with our findings in the training cohort (**Fig. 3C; Fig. S9B**). Moreover, comparable results were obtained on datasets with discontinuous developmental processes lacking transitional cells (**Fig. S10A, C**). These data distinguish CytoTRACE from RNA velocity, a recently described kinetic model that can predict future cell states, but is limited to scRNA-seq data with continuous fate transitions and genes with mRNA half-lives on the order of hours^15^ (**Fig. S10B, D**). Importantly, no significant biases in CytoTRACE performance were observed in relation to tissue type, species, the number of cells analyzed, time-series experiments versus snapshots of developmental states, or plate-based versus droplet-based technologies (**Fig. S11**).

### Differentiation-associated genes and cellular hierarchies

Given CytoTRACE’s ability to faithfully recover the direction of differentiation in nearly every evaluated dataset, we asked whether it might prove useful for discovering genetic markers of immature cells without prior knowledge of cellular phenotypes. Toward this end, we rank-ordered all genes in each benchmarking dataset based on their correlation with CytoTRACE and defined ‘ground truth’ gene sets that marked the least and most differentiated cells as annotated in the original studies. In 86% of datasets, these gene sets were significantly skewed in the correct direction toward the extreme ends of the ranked transcriptome (adjusted *P* < 0.05, GSEA^41^; **Fig. 3E; Fig. S12**). Moreover, CytoTRACE automatically prioritized well-established stem and progenitor markers, including *Kit* and *Stmn1* in the mouse bone marrow^42^ and *Axin2* and *Lgr5* in mouse intestinal crypts^43^, underscoring the utility of CytoTRACE for the *de novo* discovery of developmentally-regulated genes (**Fig. 3E**).

We next explored the potential of CytoTRACE to complement existing techniques for trajectory visualization and branch detection. By combining it with a two-dimensional force-directed layout algorithm to analyze 39,505 cells from zebrafish embryos, CytoTRACE readily revealed complex branching patterns arising during whole organism development from a fertilized egg (**Fig. S13**)^44^. Likewise, when applied to 3,427 unselected mouse bone marrow cells^42^, CytoTRACE enabled reconstruction of the directionality and lineage structure of hematopoietic development (**Fig. 4A**).

**Figure 4.**
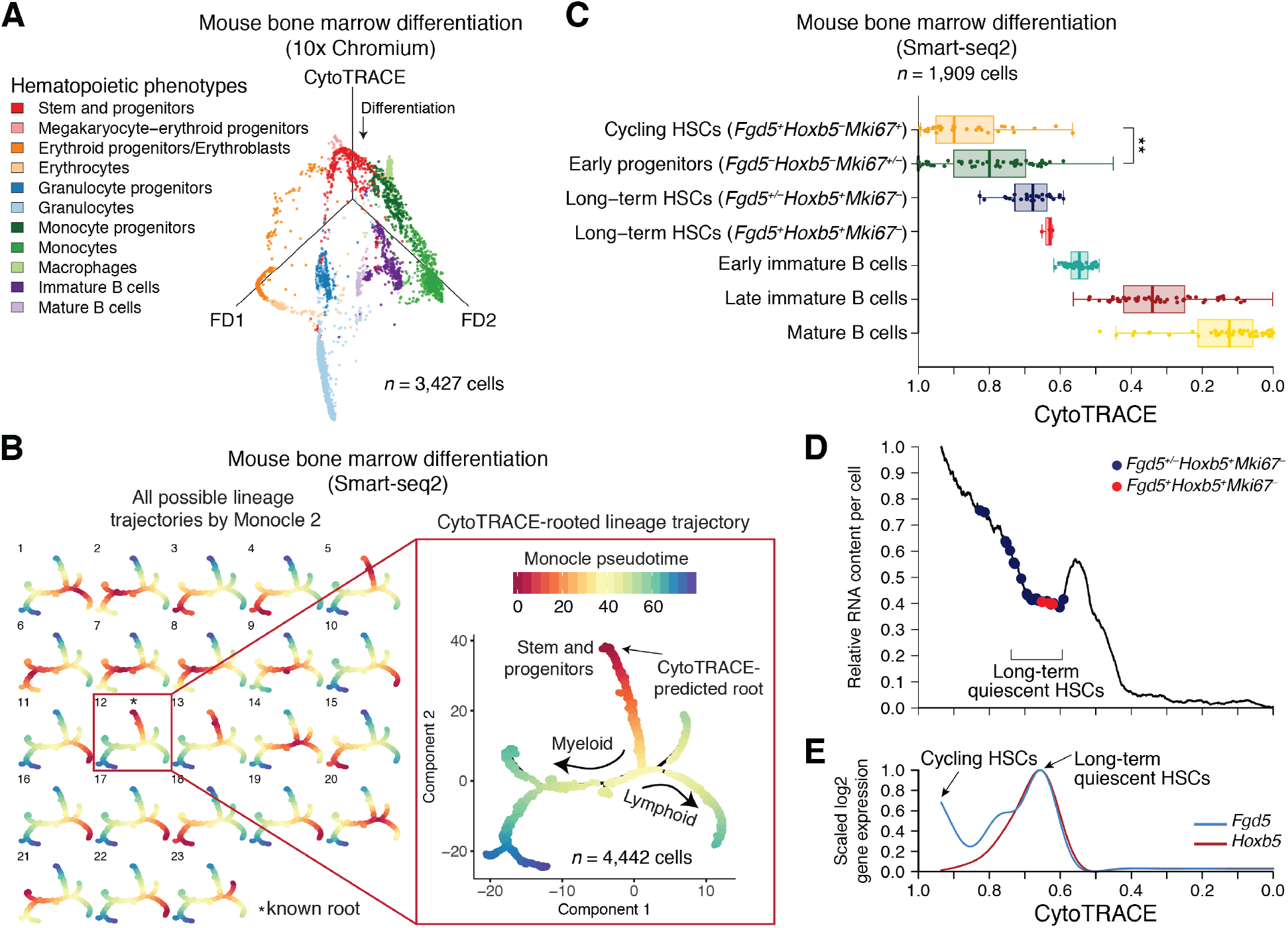
Characterization of developmental hierarchies and quiescent stem cells using CytoTRACE. (**A-E**) Application of CytoTRACE to dissect the mouse hematopoietic hierarchy by integrative analysis. (**A**) Mouse bone marrow scRNA-seq data visualized by CytoTRACE versus a force-directed layout (FD1 vs. FD2). (**B**) Combined application of CytoTRACE and Monocle 2 to delineate complex branches during mouse bone marrow differentiation without any prior information of the root. Multi-lineage tree inferred by Monocle 2 showing all 23 possible pseudotimes when the root is unknown (*left*) and automatic selection of the correct root by CytoTRACE (*right*). (**C-E**) Prioritization of quiescent and cycling hematopoietic stem cells (HSCs) in index-sorted scRNA-seq data of hematopoietic stem and progenitors (KIT^+^SCA1^+^LIN^−^) and developing B cells (TER119^−^B220^+^). All plots are identically ordered, left to right, by CytoTRACE. (**C**) Boxplots showing CytoTRACE values for candidate cycling HSCs (*n* = 31) and long-term/quiescent HSCs (*n* = 30) versus early immature B cells (*n* = 285), late immature B cells (*n* = 863), and mature B cells (n = 700). HSC subgroups were defined based on expression of *Fgd5*, a reporter gene for HSCs^67^, *Hoxb5*, a reporter gene for long-term HSCs^47^, and *Mki67*, a marker of proliferation. Although boxplots represent all analyzed cells, for clarity, a maximum of 50 cells per phenotype are displayed as points. Statistical significance was assessed by an unpaired Wilcoxon signed-rank test. \*\**P*<0.01. (**D**) Relative RNA content per cell, shown as a function of CytoTRACE (‘*Analysis of total RNA content and transcriptional diversity*’ in Methods) and displayed as the moving average of 200 cells. (**E**) Expression of *Fgd5* and *Hoxb5* ordered by CytoTRACE and displayed as a smoothing spline over the moving average of 200 cells. Data from monocytes and granulocytes (TER119^−^MAC1^+^GR1^high^) are consistent with the above results. Data in **A** and **B-E** are from ‘Bone marrow (10x)’ and ‘Bone marrow (Smart-seq2)’ datasets generated by Tabula Muris^42^, respectively (**Methods**).

Complex lineage relationships in scRNA-seq data can also be determined by dedicated branch detection tools^14^, such as Monocle 2^34^, however these approaches do not predict the starting point of the biological process. For example, when applied to 4,442 bone marrow cells^42^, Monocle 2 identified 23 possible “roots” from which to calculate pseudotime values (**Fig. 4B, left**). Only 1 of these 23 states is correct (4% of possibilities), which we define as the state that is most enriched for previously annotated stem and progenitor cells (state 12 in **Fig. 4B**). By integrating CytoTRACE with Monocle 2, the correct root was readily identified without user input (**Fig. 4B, right; Fig. S14A, B**). This facilitated identification of lineage-specific regulatory factors and marker genes during granulocyte, monocyte, and B cell differentiation (**Fig. S14C**). Similarly, when CytoTRACE and Monocle 2 were applied to 4,581 mouse intestinal cells^43^, we were able to automatically determine the root, developmental ordering, and branching processes of stem and progenitor cells differentiating into enterocyte and secretory lineages (**Fig. S14D-E**).

### Dissection of stem cell and progenitor populations

We next asked whether CytoTRACE could distinguish cycling and long-term/quiescent stem cells from their downstream progenitors^45,46^. As these populations are well-characterized in the bone marrow^3^, we investigated this question in the mouse hematopoietic system. While both cycling and quiescent hematopoietic stem cell (HSC) subpopulations^45,46^ were correctly predicted to be less differentiated, only proliferative HSCs were significantly ranked above early progenitors (**Fig. 4C**). This result was not unexpected, however, since quiescent cells have reduced metabolic activity and low RNA content^1^. By devising a simple approach to visualize inferred RNA content as a function of CytoTRACE, we observed a distinct valley in RNA abundance that coincided with elevated expression of *Hoxb5*, a recently described marker of long-term/quiescent HSCs^47^ (**Fig. 4D, E**). This analysis further confirms the value of CytoTRACE and suggests a novel approach for elucidating tissue-specific stem cells from scRNA-seq data.

### Application to neoplastic disease

Having validated CytoTRACE’s technical performance, we next applied it to a system where developmental trajectories are less well-characterized. A growing body of evidence suggests that human breast tumors are hierarchically organized and originate from subpopulations of cancer cells, called tumor-initiating cells, which are less differentiated, resistant to therapy, and implicated in relapse and metastasis^48,49^. In breast cancer, subpopulations of tumor cells within the luminal progenitor (LP) epithelium are thought to give rise to aggressive basal-like breast cancers, such as triple-negative breast cancer (TNBC)^50,51^, and possibly also to estrogen receptor positive (ER+) breast cancers^52^. However, the differentiation states and tumor-initiating properties of LP subsets remain incompletely understood^53^.

To test whether CytoTRACE can facilitate new insights into immature LP cells and their associated genes in breast cancer, we performed scRNA-seq profiling of breast tumor epithelial cells and adjacent normal epithelial cells from 8 patients with basal-like (*n* = 2) or luminal-like (*n* = 6) breast cancer. Using a Smart-seq2 protocol combined with fluorescence-activated cell sorting (FACS), we index-sorted and sequenced cells from three major human epithelial subpopulations: basal (CD49f^high^EPCAM^med-low^), luminal progenitor (CD49f^high^EPCAM^high^), and mature luminal subpopulations (ML) (CD49f^low^EPCAM^high^) (**Fig. S15A**). After removing low quality cells and applying principal component analysis to visualize the data, we confirmed three well-separated clusters of basal, LP, and ML cells, each with characteristic expression patterns of previously described lineage markers (**Fig. 5A; Fig. S15B**). No obvious clustering was observed for tumor/normal differences or by patient (**Fig. 5A**).

**Figure 5.**
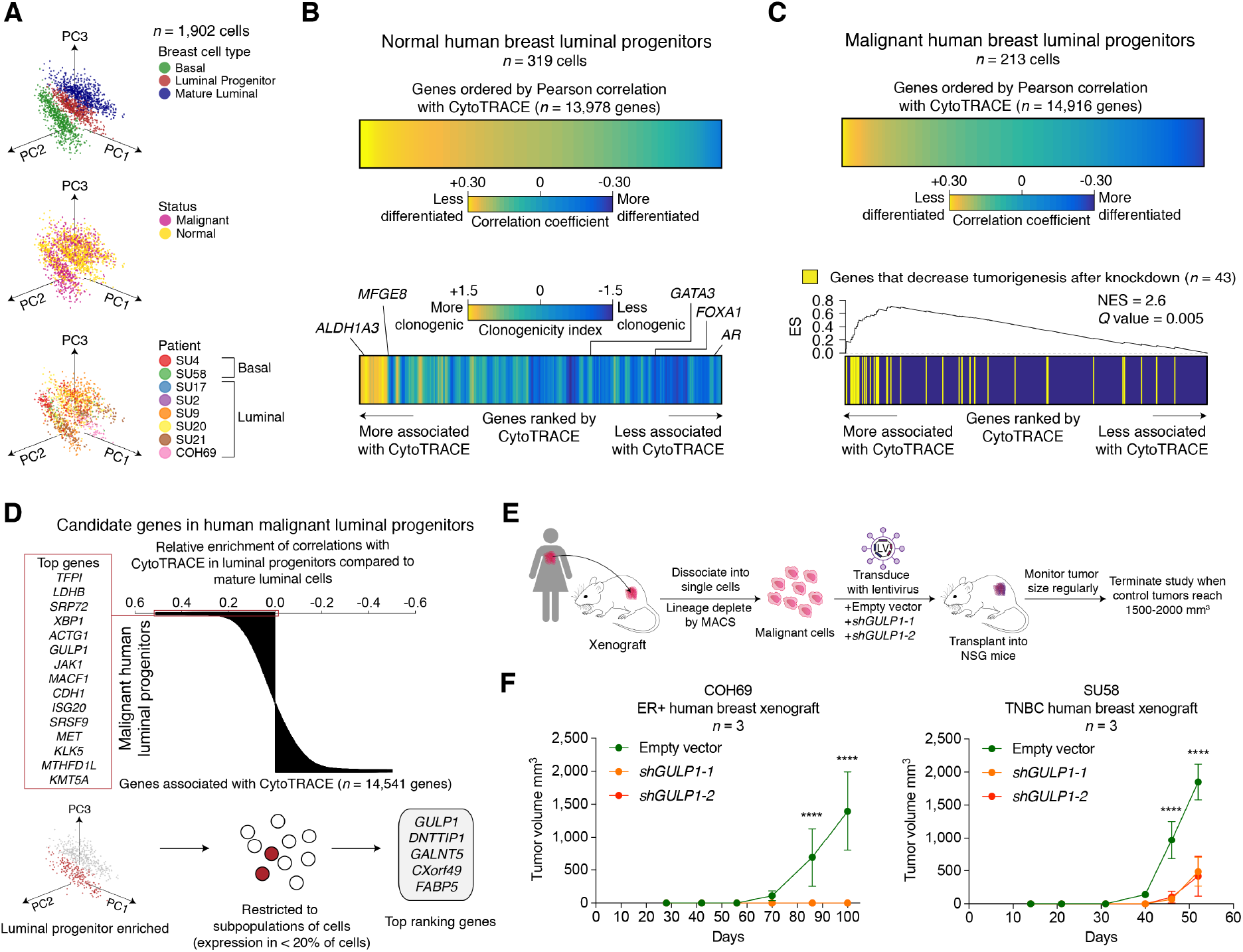
Identification of immature cell markers in normal and malignant human breast luminal progenitor cells using CytoTRACE. (**A**) Principal component analysis (PCA) plots showing scRNA-seq profiles of human basal cells (*n* = 660 cells), luminal progenitors (*n* = 532 cells) and mature luminal cells (n = 710 cells) from 8 breast cancer patients (2 basal-like and 6 luminal-like), colored by epithelial subpopulations (top), tumor vs. paired adjacent normal tissues (center), and patient (bottom). (**B**) Prediction of differentiation-associated genes in normal luminal progenitors (LPs) profiled by scRNA-seq. *Top:* Heat map showing normal LP genes ordered by their Pearson correlation with CytoTRACE. *Bottom:* Heat map depicting the association of each gene in the above plot with a ‘clonogenicity index’, defined as the log_2_-fold change in expression between highly and lowly clonogenic LPs from normal human breast^54^ (**Methods**). The clonogenicity index is displayed as a moving average of 200 genes. Key genes associated with less (*ALDH1A3, MFGE8*) and more (*GATA3, FOXA1, AR*) differentiated normal LPs are indicated. (**C**) Prediction of differentiation-associated genes in malignant luminal progenitors profiled by scRNA-seq. *Top:* Same as panel B (top) but showing genes from malignant rather than normal LPs. *Bottom:* Pre-ranked gene set enrichment analysis^68^ of 43 genes found to decrease human breast tumorigenesis in an RNAi dropout viability screen^57^ in relation to LP genes ranked by CytoTRACE (same order as above). NES, normalized enrichment score; ES, enrichment score. (**D**) Identification of candidate tumorigenic genes associated with less differentiated malignant human LPs. *Top:* Genes rank-ordered by the difference in their correlations with CytoTRACE in malignant LPs versus malignant mature luminal cells (MLs). The top 15 genes that are predicted to be specifically associated with less differentiated LPs are indicated (left). *Bottom:* Schema for the identification of genes that are ranked as above, but that are also more highly expressed in malignant LPs than MLs (log_2_ fold change > 0; Benjamini-Hochberg adjusted *P* < 0.05, unpaired two-sided *t*-test) and that are expressed by a subpopulation of LPs (<20% of cells). The top 5 filtered genes are shown (right). (**E**) Schema for shRNA knockdown of *GULP1* in a human breast cancer xenograft model. Lineage-depleted breast cancer epithelia cells from patient-derived xenografts were transduced with lentivirus containing either an empty vector control or shRNA targeting *GULP1* and transplanted into immunodeficient NSG mice in triplicates. Tumors were monitored weekly until control tumors reached a size of 1500-2000 mm^3^. (**F**) Growth of human breast cancer xenografts from two patients, one with ER+ luminal-type cancer (left) and one with triplenegative breast cancer (right) after lentiviral transduction with empty vector or shRNA targeting *GULP1*. Mean tumor volume with 95% confidence interval is shown for 6 time points (*n* = 3 mice). **** *P* < 0.0001.

To validate the ability of CytoTRACE to define LP differentiation states, we started by rank-ordering genes in adjacent normal LPs by their correlation with CytoTRACE (**Fig. 5B, top**). We found that previously described marker genes of less differentiated normal LPs (*ALDH1A3* and *MFGE8*)^54,55^ and more differentiated normal LPs (*GATA3, FOXA1*, and *AR*^54,56^) were successfully enriched by this approach (**Fig. 5B, bottom**). Moreover, genes that were upregulated in highly clonogenic normal LPs^54^ were skewed toward genes predicted to mark less differentiated cells (**Fig. 5B, bottom**).

Given these favorable results, we next sought to identify LP genes associated with tumorigenesis. By rank-ordering LP genes in malignant cells by their correlation with CytoTRACE (**Fig. 5C, top**), we observed a highly significant enrichment of genes whose knockdown by RNA interference (RNAi) led to decreased viability of tumor cells in patient-derived xenograft (PDX) models of TNBC^57^ (*Q* = 0.002, GSEA; **Fig. 5C, bottom; Fig. S16**). Moreover, when we applied CytoTRACE to prioritize genes in tumor LPs as compared to tumor MLs, the latter of which are developmentally downstream of LPs in normal breast^54,58^, the top 15 genes included known members of tumorigenic pathways in breast cancer (e.g., *MET, JAK1*, and *XBP1*^59–61^), as well as novel candidates (e.g., *GULP1*) (**Fig. 5D, top**). To further refine this list, we focused on genes that were (1) more highly expressed in tumor LPs than MLs and (2) expressed in a subpopulation of tumor LPs (<20% of cells) (**Fig. 5D, bottom**). After applying this filter, *GULP1* emerged as the top candidate (**Fig. 5D, bottom right**).

GULP1 is an engulfment adaptor protein and is the human homolog of Drosophila Ced6^62^. Moreover, the murine homolog of GULP1 is a specific marker of mouse HSCs, suggesting a possibly conserved role of this gene in other immature cell states (**Fig. S17A**). Since the role of GULP1 in human breast cancer has not been established, we measured the effect of *GULP1* knockdown on the proliferation of metastatic TNBC cell lines, MDA-MB-231 and MDA-MB-468, compared to an empty vector control (**Fig. S17B-E**). *GULP1* knockdown significantly reduced proliferation of MDA-MB-231 and MDA-MB-468 as measured by a colorimetric assay for metabolic activity (**Fig. S17E**). Next, we tested the effect of *GULP1* knockdown in PDXs from ER+ and TNBC patients from our single-cell cohort (**Fig. 5E**). We found that knockdown of *GULP1* significantly reduced tumor growth in the TNBC sample and completely abrogated the ER+ tumor compared to empty vector control (**Fig. 5F**).

Taken together, these data suggest a novel role for *GULP1* in human breast cancer tumorigenesis and demonstrate the promise of CytoTRACE for the discovery of malignant cell differentiation states and new therapeutic targets.

## Discussion

Efforts to characterize single-cell transcriptomes in diverse tissues, organs, and whole organisms^42,63^ have underscored the need for robust RNA-based determinants of developmental potential. In our analysis of ~19,000 features across 42 developmental processes and nearly 150,000 single cells, we found that gene counts, or the number of detectably expressed genes per cell, powerfully associates with transcriptional diversity and differentiation status. Although anecdotally observed in specific experimental systems (mouse alveolar epithelial development, zebrafish thrombopoiesis, and neuron differentiation from hESCs^26–28^), we demonstrate for the first time that this association (1) outperforms most stemness inference tools and pre-defined molecular signatures from a compendium of nearly 19,000 RNA-based features, (2) is generally independent of species, platform, and tissue type, and (3) is broadly applicable throughout cellular ontogenesis.

Although previous studies have demonstrated a global reduction in chromatin accessibility and/or plasticity during lineage commitment in specific developmental settings (e.g. embryonic stem cells, intestinal stem cells, and neural stem cells^35,36,38,64^), our quantitative findings extend the scope of this result. Moreover, as has been previously shown^65^, our data indicate that variability in gene counts between phenotypically identical single cells is not exclusively due to drop-out events, but also due to differential sampling of the transcriptome (**Fig. S3**). Our data are therefore consistent with a model in which less mature cells maintain looser chromatin to permit wider sampling of the transcriptome, while more differentiated cells generally restrict chromatin accessibility and transcriptional diversity as they specialize (**Fig. S6C**)^66^. Future studies will be needed to further confirm the validity of this model and assess its relevance across diverse tissue compartments, developmental time points, and phenotypic states.

The identification of gene counts as a leading measure of cellular differentiation status motivated the development of CytoTRACE, a computational framework that leverages and significantly improves upon gene counts for resolving differentiation states at the single-cell level. Unlike most existing methods for lineage trajectory analysis^8–15^, however, CytoTRACE can predict both the relative state and direction of differentiation in a manner that is independent of specific timescales or the presence of continuous developmental processes in the data. CytoTRACE is also agnostic to tissue type, species, and scRNA-seq platform.

We anticipate that these advantages will enable significant applications. For example, by using CytoTRACE to analyze scRNA-seq profiles of human breast tumors, we identified new candidate genes associated with less-differentiated luminal progenitor cells and established a novel role for *GULP1* in breast tumorigenesis. These data emphasize the utility of CytoTRACE for characterizing tumor differentiation hierarchies and for discovering novel biomarkers and therapeutic targets. Furthermore, by integrating RNA content with CytoTRACE, we demonstrated, for the first time to our knowledge, that quiescent adult stem cells can be distinguished from downstream progenitors using an unsupervised *in silico* approach. Given the immense regenerative potential of quiescent stem cells, their identification in human tissues has broad implications in regenerative medicine and cancer treatment.

While CytoTRACE can recapitulate developmental orderings from single lineages to whole organisms, several challenges remain. For example, although the direction of differentiation was predicted correctly in nearly all datasets, 12% of cases were mischaracterized. These datasets also proved problematic for other approaches, suggesting there may be opportunities for future enhancements. In addition, CytoTRACE is currently expressed in rank space, which is not directly comparable across different datasets. Efforts to overcome these challenges are underway.

In summary, we conclude that the number of expressed genes per cell is a hallmark of developmental potential. By exploiting this data-driven property of scRNA-seq data, we developed a broadly applicable framework for resolving single-cell differentiation hierarchies. We envision that our approach will complement existing scRNA-seq analysis strategies, with implications for the identification of immature cells and their developmental trajectories in complex tissues throughout multicellular life.

## Acknowledgments

We thank A. Chaudhuri, A. Gentles, and A. Alizadeh for critical feedback on the manuscript. We are grateful to S. Bobo for assistance with patient specimen acquisition, R. Sinha and C.K.F. Chan for provision of data and resources, C.L. Liu for assistance with the website, P. Lovelace and S. Weber for their support and assistance with FACS, and S. Sim for assistance with processing and sequencing scRNA-seq libraries. This work was supported by grants from the National Cancer Institute (A.M.N., R00CA187192-03; M.F.C., 5R01CA100225-09; G.S.G., PHS Grant Number CA09302), the Stinehart-Reed foundation (A.M.N.), the Stanford Bio-X Interdisciplinary Initiatives Seed Grants Program (IIP) (A.M.N., M.F.C), the Virginia and D.K. Ludwig Fund for Cancer Research (A.M.N., M.F.C), the U.S. Department of Defense (M.F.C., W81XWH-11-1-0287 and W81XWH-13-1-0281; S.S.S., W81XWH-12-1-0020), a National Science Foundation Graduate Research Fellowship (DGE-1656518 to M.J.B.), Stanford Bio-X Bowes Graduate Student Fellowship (G.S.G.), and the Stanford Medical Science Training Program (G.S.G.).

## Author contributions

G.S.G. and A.M.N. developed the concept for CytoTRACE, designed related experiments, and analyzed the data with assistance from S.S.S., D.J.W., and M.F.C. G.S.G. and A.M.N. wrote the manuscript with assistance from S.S.S. G.S.G. and A.M.N. performed the bioinformatics analyses with assistance from D.J.W., A.B., A.M., M.J.B., and F.L. A.M. and A.B. designed the website with input from G.S.G. and A.M.N. S.S.S. generated the human breast cancer singlecell RNA-sequencing data with assistance from A.H.K., R.W.H., S.C., M.Z., F.A.S., N.A.L., D.Q., and F.B.Y. S.S.S. performed the mouse experiments under the supervision of M.F.C. F.D. assisted with the collection of patient specimens. All authors commented on the manuscript at all stages.

## Data and materials availability

Details of publicly available datasets are provided in **Methods**. Single-cell RNA-seq expression data generated in this study are hosted at https://cytotrace.stanford.edu with a GEO accession number pending.

