## Supplementary Materials for "Single-cell transcriptional diversity is a hallmark of developmental potential"

#### Table of Contents:

##### Methods

1. External datasets
2. *In silico* screen for features associated with differentiation
3. Stemness inference tools
4. Performance assessment and benchmarking
5. Analysis of gene counts in non-cycling and cycling cells
6. Analysis of gene counts and pluripotency genes during human embryonic development
7. Analysis of gene counts across mouse cellular ontogeny
8. Analysis of gene counts in pooled transcriptomes and following imputation in single-cell transcriptomes
9. Dependency of gene counts on the number of reads per cell
10. Analysis of total RNA content and transcriptional diversity
11. Chromatin accessibility analysis
12. Overview of CytoTRACE
13. Analysis of differentiation-associated genes
14. 3D visualization of differentiation trajectories with CytoTRACE
15. Combined root selection and branch detection with CytoTRACE and Monocle 2
16. RNA velocity
17. Human subjects
18. Tissue dissociation
19. Flow cytometry
20. Single-cell expression profiling of human breast tumors
21. Transcript quantification and cluster analysis of breast tumors
22. Clonogenicity index
23. RNAi dropout viability screen
24. Lentivirus production
25. Xenograft tumor cell transduction and engraftment
26. Cell viability assay
27. Statistical analysis
28. Software implementation and website
29. Code availability
30. Data availability

##### Supplementary Figures

Figure S1 *In silico* screen for features of differentiation

| | |
|---|---|
| Figure S2 | Association between gene counts and differentiation in cycling and non-cycling cells |
| Figure S3 | Approaches to overcome sparsity in scRNA-seq data degrade the predictive performance of gene counts |
| Figure S4 | Impact of the number of reads per cell on gene counts |
| Figure S5 | Impact of different minimum expression thresholds on the association between gene counts and developmental potential |
| Figure S6 | Association between gene counts and RNA abundance |
| Figure S7 | Dynamics of gene counts, transcriptional diversity, and global chromatin accessibility during *in vitro* differentiation of hESCs into lateral mesoderm |
| Figure S8 | Development and robustness of CytoTRACE |
| Figure S9 | Prediction of single-cell differentiation states in 33 validation datasets |
| Figure S10 | Prediction of single-cell differentiation states without transitional cells |
| Figure S11 | Robustness of CytoTRACE to variation in dataset characteristics |
| Figure S12 | Evaluation of CytoTRACE for prioritizing developmental marker genes |
| Figure S13 | Reconstruction of early zebrafish development |
| Figure S14 | Utility of combining CytoTRACE with Monocle 2 to identify rooted developmental hierarchies and lineage-specific genes without prior knowledge |
| Figure S15 | Gating scheme and marker validation of epithelial subpopulations from human breast tumors and adjacent normal tissues |
| Figure S16 | Validation of CytoTRACE for prioritizing markers of breast tumorigenesis using results from an RNAi dropout viability screen |
| Figure S17 | Enrichment of *Gulp1* in hematopoietic stem cells and validation of *GULP1* knockdown |

### Methods

1. External datasets

We selected 42 scRNA-seq datasets from 34 studies for the identification and validation of RNA-based correlates of developmental potential (see *'Data availability'*). The discovery (or training) cohort (*n* = 9 datasets) was curated to prioritize datasets previously used for benchmarking published stemness inference tools[12,18-20,29], and to include representative datasets of multi-branching hematopoiesis[30] and neuronal differentiation[26]. The validation cohort (*n* = 33 datasets) was curated to include a broad array of differentiation trajectories, including 17 datasets from a recent benchmarking study of lineage trajectory inference methods[14]. For stringency, we restricted our analysis to datasets with experimentally-confirmed differentiation states and *in vivo* time-series experiments. Datasets were excluded if they (1) did not directly capture differences in developmental potential (e.g., cell activation or polarization, metabolic shifts, and response to stimuli were omitted), (2) had clear technical batches with highly uneven representation of cellular phenotypes between batches, (3) were previously filtered to represent only a small subset of the transcriptome, or (4) captured an *in vitro* time-series experiment of a heterogeneous population of cells without prior annotations of cellular phenotypes. Finally, we limited our compendium to non-redundant differentiation or developmental processes with the exception of those from different species or platforms.

Pre-processed gene expression matrices were downloaded as provided by the authors, and were rescaled to transcripts per million (TPM) or counts per million (CPM) as appropriate. Cells with fewer than 10 detectable genes or no detectable genes among the top 1,000 most dispersed genes (calculation described in '*Overview of CytoTRACE*') were removed. For datasets that clustered by technical batches ('Aging HSCs (Smart-seq2)'[69] and 'Lung fibroblast (C1)'[70]), batch correction was applied with ComBat[71,72], as implemented in *sva* v3.24.4 (R package). Batch correction improved CytoTRACE performance in the 'Lung Fibroblast (C1)'[70] dataset from a Spearman rho of 0.38 before correction to 0.79 after correction. For 'Intestine (Drop-seq)'[43], batches 'B1' and 'B2' out of 10 batches annotated by the authors clustered separately and were therefore excluded from further analysis. Of note, *homologene* v1.1.68 (R package), which references HomoloGene v68 (NCBI), was applied to reconcile species-specific differences in gene symbols between evaluated features and datasets by converting gene symbols in the latter. For single-cell expression data derived from *Macaca fascicularis*[73], we used gene symbols from the closest relative of *M. fascicularis* in HomoloGene v68, *Macaca mulatta*. Analyses requiring human or mouse-specific gene symbols were not performed on a recently published dataset of whole planaria (*Schmidtea mediterranea*)[74] owing to the absence of this organism from HomoloGene v68. Additional pre-processing steps for specific datasets are described below.

- 'Aging HSCs (Smart-seq2)'[69]: Long-term hematopoietic stem cells (LT-HSCs), short-term hematopoietic stem cells (ST-HSCs), and multipotent progenitors (MPPs) from old and young mice were combined into one table to include all single cells from the study without introducing redundancy in the validation cohort.
- 'Dentate gyrus phenotypes (10x)'[33], 'Dentate gyrus timepoints (10x)'[33], 'Peripheral blood (10x)'[75], 'Neural stem cells (Drop-seq)'[76], 'Whole planaria (Drop-seq)'[74], 'Thymus (Drop-seq)'[77], and 'Early zebrafish (Drop-seq)'[44]: To accommodate memory limitations of several lineage trajectory inference methods, these larger datasets were randomly down-sampled to 6,000 single cells prior to analysis (except for 'Peripheral blood (10x)'[75], which was down-sampled to 5,000 single cells). Of note, CytoTRACE was also run on all cells without down-sampling, achieving comparable performance to the down-sampled data (data not shown).

- ‘Pre-implant human embryo (Tang et al.)’[20,78]: As previously described[20], two cells from the morula stage, ‘Morula #1 – Cell #3’, ‘Morula #1 – Cell #8’, were excluded as outliers since they clustered with cells from the late blastocyst stage.
- ‘Human skeletal muscle myoblasts, or HSMM (C1)’[29]: As previously described[29], interstitial mesenchymal cells were considered contaminants and excluded.
- ‘Neural stem cells (Drop-seq)’[76]: As previously described[76], E15.5 neural stem cells were acquired from a different technical batch and therefore excluded.

In each dataset, only phenotypes involved in a known differentiation process described by the authors were included.

2. *In silico* screen for features associated with differentiation

To identify RNA-based factors correlated with cellular potency, we screened 18,711 candidate features in a training cohort comprising nine scRNA-seq datasets (‘*External datasets*’). These features included: (1) all 17,810 available gene sets from the Molecular Signatures Database (v6.2), spanning hallmark, positional, curated, motif, computational, gene ontology, oncogenic, and immunogenic gene sets[23,79,80], (2) 896 gene sets covering experimentally determined transcription factor binding sites from the Encyclopedia of DNA Elements (ENCODE)[24] and ChIP Enrichment Analysis (ChEA)[25], (3) stemness inference tools based on models of transcriptional entropy (StemID[18], SCENT[19], and SLICE[20]), (4) a one-class logistic regression model (mRNA stemness index; mRNAsi)[22], and (5) gene counts, defined as the number of transcripts per cell >0 (TPM datasets) or the number of counts/UMIs per cell >0 (CPM datasets). Gene set features were calculated using single-sample gene set enrichment analysis (ssGSEA) as implemented in the R GSVA package (v1.30.0)[81]. Entropy measures and mRNAsi were calculated as described below (‘*Previous methods*’).

3. Stemness inference tools

The following stemness inference tools[18-20] were applied using default settings unless stated otherwise. R packages: *StemID* v0.0.0.9000 (entropy function only)[18], *scent* v1.0[19], and *SLICE* v0.99.0[20]. For *SCENT*, the default human protein-protein interaction network provided by the authors was used to calculate entropy. For *SLICE*, scRNA-seq datasets derived from primates (human and macaque) were analyzed with the human gene-gene Kappa similarity matrix whereas scRNA-seq datasets from mouse and zebrafish were analyzed with the mouse similarity matrix. mRNAsi[22] was trained and run per the online instructions to replicate the model employed in the original study (http://tcgabiolinks.fmrp.usp.br/PanCanStem/mRNAsi.html). For methods requiring $\log_2$ expression data (all but StemID), a pseudo-count of 1 was added prior to $\log_2$ adjustment. The *homologene* v1.1.68 R package, which references HomoloGene v68 (NCBI), was used to convert gene symbols in scRNA-seq data matrices in cases where a different species was expected: SCENT (human), SLICE (human or mouse), mRNAsi (human), or ssGSEA (human) (for further details, see ‘*External datasets*’ above).

4. Performance assessment and benchmarking

To numerically represent differentiation status, each author-supplied cellular phenotype was assigned a rank based on its experimentally-confirmed differentiation or developmental status, where the minimum rank corresponded to the least mature cell and the maximum rank to the most mature cell (**Fig. 1B**, **bottom right**). To evaluate performance at the phenotype level, the output from each feature or method was mean-aggregated by phenotype, ranked, and compared to ground truth ranks by Spearman correlation. To assess performance at the single-cell level, the results of each feature/method were weighted by the total number of cells per phenotype, ranked, and compared against ground truth ranks by Spearman correlation using *weights*[82] v0.90 (R package). Mean-aggregated or single-cell values were ranked in ascending order, such that higher values corresponded to lower ranks. Thus, features and methods that decreased with differentiation yielded positive correlations, and vice versa. For datasets with multiple lineages (i.e., 'Dentate gyrus phenotype (10x)'[33], 'Bone marrow (10x)'[42], 'Bone marrow (Smart-seq2)'[42], 'Mesoderm (C1)'[32], 'Lung fibroblast (C1)'[70]), we considered each lineage as a separate differentiation process, and calculated performance as the mean of the lineage-specific Spearman correlation coefficients in each dataset.

5. Analysis of gene counts in non-cycling and cycling cells

The number of unique transcripts expressed by a cell can be influenced by multiple factors, including metabolic activity, secretory function, and cell cycle status[83]. To measure the effect of cell cycle status on the association of gene counts with differentiation, we stratified single cells in each dataset into 'cycling' and 'non-cycling' groups based on their expression of cell cycle genes. Cell cycle genes were selected from the 'GO_CELL_CYCLE' (GO: 0007049) gene set, and expression was calculated as the geometric mean of the gene set for each single cell. To stratify single cells into 'cycling' and 'non-cycling', we fit a two-component Gaussian mixture model to each dataset in the training cohort (*mclust* v5.3, R package), and defined the threshold as the expression value at which a cell has an equal probability of being classified as 'cycling' and 'non-cycling' (posterior probability = 0.5). Bimodality was tested by performing a Chi-square test on the difference of the log likelihoods of a two-component Gaussian mixture model versus a single-component Gaussian mixture model. Datasets with insufficient evidence of bimodality (Chi-square $P \geq 0.05$), and datasets with fewer than 10 cells assigned to 'cycling' or 'non-cycling' were excluded from the analysis. For each qualified dataset in the training cohort ($n = 5$, i.e. 'HSPCs (C1)', 'hESC *in vitro* (C1)', 'HSMM (C1)', 'Lung development (C1)', 'AT2/AT1 lineage (C1)', and '*in vitro* NPCs (C1)'; see *'Data availability'*), performance was calculated as described above.

6. Analysis of gene counts and pluripotency genes during human embryonic development

For the analysis in **Figure 1D**, we utilized single cell transcriptomes of 12 experimentally-validated, purified cell phenotypes from two *in vitro* plate-based datasets, 'Pre-implant human embryo (Tang et al.)' and 'Mesoderm (C1)' (see *'Data availability'*). The ground truth order for each phenotype was determined by first binning it into a known potency group and then ordering based on definitions provided by authors of the studies or experimentally validated elsewhere. These phenotypes, from least to most differentiated with ranked developmental stage indicated in parenthesis, included, zygote[78] (1), 2-cell[78] (2), 4-cell[78] (3), 8-cell[78] (4), morulae[78] (5), human embryonic stem cell[32] (6), anterior primitive streak[32] (7), paraxial mesoderm[32] (8), somitomere[32] (9), early somite[32] (10), dermomyotome[32] (11), and sclerotome[32] (11). The late blastocyst[78] from 'Pre-implant human embryo (Tang et al.)' (see *'Data availability'*) was omitted as it includes heterogenous cell types with different developmental potencies. To fairly compare across different datasets with varying sequencing depths, raw reads were acquired for each dataset from the Sequence Read Archive and aligned to the GENCODE v25 reference transcripts (GRCh38.p7) using *Salmon*[84] v0.7.2 with flags -l IU for paired-end reads, -l U for single-end reads, and --seq-bias for both. All single-cell transcriptomes were down-sampled to 1 million mapped reads prior to analysis. The number of genes per cell (gene counts) and ssGSEA[81] of pluripotency genes[21] were calculated as described above (see '*In silico screen for features associated with differentiation*').

7. Analysis of gene counts across mouse cellular ontogeny

We restricted our analysis in **Figure 1E** to plate-based scRNA-seq experiments capturing *in vivo* developmental processes in mice. These criteria were selected owing to the superior capture rates of plate-seq technologies[85], and the availability of mouse datasets covering *in vivo* processes with well-defined phenotypes across all potency levels[31]. *In vivo* processes were prioritized to avoid alterations in gene expression induced by *in vitro* systems. The ground truth order for each phenotype was determined by first binning it into a known potency group and then ordering based on definitions provided by authors of the studies or experimentally validated elsewhere. These phenotypes, from least to most differentiated with ranked developmental stage indicated in parenthesis, included, zygote[86] (1), 2-cell[86] (2), 4-cell[86] (3), 8-cell[86] (4), 16-cell[86] (5), early blastocyst[86] (6), middle blastocyst[86] (7), late blastocyst[86] (8), hematopoietic stem cell progenitor (HSCP)[30] (9), hematopoietic stem and progenitors (KLS)[42] (10), multi-lineage primed hematopoietic progenitor[30] (10), embryonic bipotent alveolar progenitor[27] (11), monocyte-dendritic cell progenitor[30] (12), monocyte-dendritic cell progenitor[87] (12), type II pneumocyte (AT2)[27] (13), common dendritic progenitor[87] (13), myelocyte[30] (14), type I pneumocyte (AT1)[27] (15), erythroid progenitor[30] (15), granulocyte progenitor[30] (15), early immature B cell[42] (15), megakaryocyte progenitor[30] (15), monocyte progenitor[30] (15), monocyte progenitor and monocytes[42] (15), pre-dendritic cell[87] (15), late immature B cell[42] (16), early immature (proliferating) granulocytes[42] (16), mature B cell[42] (17), late immature granulocytes[42] (17), and mature (resting) granulocytes[42] (17). To fairly compare across different datasets with varying sequencing depths, raw reads were acquired for each dataset from the Sequence Read Archive and aligned to the GENCODE vM15 reference transcripts (GRCm38.p5) using *Salmon*[84] v0.7.2 with flags -l IU for paired-end reads, -l U for single-end reads, and --seq-bias for both. For all cells with greater than 1 million mapped reads ($n$ = 3,875), single cell transcriptomes were down-sampled to 1 million reads and gene counts were recalculated as described below '*Dependency of gene counts on the number of reads per cell*'. For cells with fewer than 1 million mapped reads ($n$ = 1,184), a linear regression model was learned from the relationship between gene counts and the log number of reads at down-samples of 10%, 20%, 30%, 40%, 50%, 60%, 70%, 80%, and 90% and used to predict the expected gene counts at 1 million reads (mean $R^2$ = 0.997 ± 0.002).

8. Analysis of gene counts in pooled transcriptomes and following imputation in single-cell transcriptomes

For the analysis presented in **Figure S3A, B, E**, FASTQ files from bulk RNA-seq and scRNA-seq analyses were downloaded from SRA (SRP073808) and aligned to GENCODE v25 reference transcripts (GRCh38.p7) as described in '*Transcript quantification and cluster analysis of human breast tumors*'. For **Figure S3C**, we created pseudo-bulk mixtures for each dataset from the training cohort ($n$ = 9) by pooling reads across all single cells of the same phenotype. Separately, to assess the impact of single-cell dropout imputation on gene counts (**Fig. S3D**), we applied *scImpute*[33] v0.0.8 (R package) to each dataset in the training cohort. The *scImpute* 'Kcluster' parameter, which requires input of the number of clusters by the user, was set to the number of unique phenotypes defined by the original authors of each dataset. Otherwise, default settings were used. For each analysis above, we determined gene counts as the total number of detectably expressed genes; Spearman correlation (weighted by number of cells per phenotype for single cell correlations) between gene counts and known differentiation status was used to evaluate performance.

To visualize heterogeneity in detectably expressed genes across single cells (**Fig. S3E**), we applied the 'oncoPrint' function from *ComplexHeatmap*[88] v1.20.0 (R package) to 67 single-cell transcriptome profiles of dermomyotome cells[32]. Gene expression was binarized into 'expressed' (TPM > 0) and 'not detected' (TPM = 0) prior to analysis. Gene counts from single-cell transcriptomes were compared against pooled single-cell transcriptomes (*n* = 67) and a bulk RNA-seq profile of the same cell type (~50,000 cells, K.M. Loh, personal communication).

9. Dependency of gene counts on the number of reads per cell

To assess the relationship between gene counts and the number of reads per cell, we selected seven count-based scRNA-seq datasets from the training cohort with available phenotypic annotations from the Sequence Read Archive (NCBI). Raw FASTQ files were downloaded and aligned to either GENCODE v25 reference transcripts (GRCh38.p7; human only) or GENCODE v15 reference transcripts (GRCm38.p5; mouse only) as described above (see '*Transcript quantification and cluster analysis of human breast tumors*'). After transcript quantification, the total number of reads per cell was down-sampled to 50%, 10%, or 1% of the original library size, or 500,000, 100,000, or 10,000 reads per cell. To maintain the original distribution of reads/cell, we randomly sampled each gene according to its probability of expression (e.g., the number of reads assigned to each gene *i* in cell *j* divided by the total number of reads assigned to cell *j*) until the desired number of reads was achieved. We then recalculated gene counts on each down-sampled transcriptome.

10. Analysis of total RNA content and transcriptional diversity

To evaluate the relationship between gene counts and total RNA content, we analyzed the number of unique molecular identifiers (UMIs), the total number of reads after ERCC spike-in normalization, and the total transcript abundance following cell lysis as inferred by Census[34]. For datasets with UMIs (**Fig. S6A**), the total number of UMIs was calculated as the sum of all UMIs per cell. For datasets with ERCCs (**Fig. S6B**), transcript abundance was inferred from the 92 ERCC spike-in standards (Thermo Fisher) using the linear regression model contained in the 'relative2abs' function in the *monocle* v2.10.1 R package. For **Figure S6C**, transcript abundance was inferred with Census using the 'relative2abs' function in *monocle* v2.10.1 R package with default settings[9]. For both ERCC- and Census-based inference of RNA content, datasets were TPM or CPM normalized prior to input and total RNA content was quantified as the sum of the inferred counts after ERCC or Census normalization. For **Figure 2A**, single cells from 'Mesoderm (C1)' (see *'Data availability'*) were reanalyzed as described above in "*Analysis of gene counts in pooled transcriptomes and following imputation in single-cell transcriptomes*". Spliced transcript abundances were calculated as the total number of detectable (>0 reads) protein-coding transcripts per cell based on GENCODE v25 reference transcript annotations (GRCh38.p7).

11. Chromatin accessibility analysis

For the analysis presented in **Figure 2B, C** and **Figure S7A, B**, ATAC-seq reads were downloaded from the Sequence Read Archive (NCBI)[32], aligned to the hg38 reference genome (GRCh38, version GCA_000001405.15) with *bwa-mem*[89] v0.7.17(r1188), and filtered for quality (-q 30 –f 0x2) and duplicates ('MarkDuplicates' tool in *Picard*[90] v2.17.8). Reads aligned to scaffolds and chrY or chrM were discarded. Reads were shifted (+4/-5) to reflect true Tn5 insertion sites. For each biological replicate (reads from technical replicates were pooled), peaks were called using *MACS2*[91] v2.1.1 (--call-summits --nomodel --shift -100 --extsize 200 -p 0.1 –keep-dup all). To generate a master atlas of most reproducible peaks, an IDR <0.05 filter was applied to pairwise combinations of biological replicates. Resulting peaks from all replicates and cell types were pooled, and overlapping peaks were merged using the ‘merge’ tool in *bedtools*[92] v2.25.0. This approach yielded a total of $n$ = 226,166 peaks.

For peak shape visualization, peaks were centered to their summit (defined as the base with maximum coverage) and adjusted to a width of 1kb. For each peak, 100 bins of 10 bp width were generated, and per-bin coverage was calculated after centering reads around the tn5 insertion site and extending to 200 bp. Peaks were then re-centered to the bin with maximum coverage, followed by CPM normalization of counts per replicate. CPM values were averaged across biological replicates, $\log_2$ transformed, and sorted by total signal per peak. For plotting, values were capped at 0.2 CPM.

To quantify the number of accessible regions in each cell type, a set of background peaks was generated by randomly sampling locations across the genome, while maintaining the number and distribution of peak lengths per chromosome from our master peak atlas. This was done by randomizing peak sites in the master peak atlas using the ‘shuffle’ tool in *bedtools*[92]. Overlap with self and with called peak regions was disallowed (-chrom, -excl and -noOverlapping options). Signal at background regions was calculated as described above. Values were CPM normalized in combination with master atlas peaks, and per-cell type background thresholds were defined as mean summit values across background regions. Master atlas peaks were considered accessible in a given cell type if the summit signal exceeded a global background threshold defined as the mean of all per-cell type background thresholds.

12. Overview of CytoTRACE

CytoTRACE is a computational framework that leverages single-cell gene counts, covariant gene expression, and local neighborhoods of transcriptionally similar cells to predict ordered differentiation states from scRNA-seq data. The approach consists of several key steps, each of which is schematically depicted in **Figure 3A** and described below.

*Gene counts signature.*
As single-cell RNA-sequencing was designed to capture gene expression, not gene counts, we reasoned that genes whose expression patterns correlate with gene counts might better predict the ordering of single cells by differentiation status. Indeed, a gene counts signature (GCS), defined as the geometric mean of genes that are most correlated with gene counts, outperformed all evaluated features in the training cohort (**Figs. S1B and S8A**). Although GCS was robust across a broad range of gene set sizes in the training cohort ($n$ = 5 to 1,000 genes), the top 200 genes maximized performance, albeit only slightly (**Fig. S8C**). We used the top 200 genes throughout this work and as input to CytoTRACE.

*Census.*
Previous studies have demonstrated that relative transcript counts, or the estimated abundances of mRNA molecules in the cell lysate, can improve the detection of differentially expressed genes over data represented as read counts (e.g., TPM)[34,93]. Relative transcript counts can be obtained with UMIs or exogenous RNA standards (e.g. ERCC), but not all platforms use UMIs (e.g., plate-based methods such as Smart-seq2), and spike-ins are not always added. Census is an algorithm that infers relative transcript counts from read counts without UMIs or ‘spike-in’ standards[34]. In our study, we found a nearly identical relationship between gene counts per single cell and the transcript counts per single cell inferred by Census ($R^2 \approx 1$; **Fig. S6C**). Given the ease of calculating gene counts and the advantages of Census transformation[34], we rescaled each single-cell transcriptome to $G_j$, the total number of detectable genes in cell $j$, as follows:

$$\mathbf{X}^*_{ij} = G_j \cdot \frac{\mathbf{X}_{ij}}{\sum_{q=1}^{n} \mathbf{X}_{qj}}, \forall i \in \{1, \dots, n\}, \forall j \in \{1, \dots, k\}$$

where $\mathbf{X}$ is an $n \times k$ matrix with $n$ genes and $k$ single-cell transcriptomes in non-log linear space, and $\mathbf{X}_{ij}$ is the quantity of transcripts or reads assigned to gene $i$ in cell $j$ for TPM and CPM matrices, respectively. The resulting gene expression matrix $\mathbf{X}^*$ was $\log_2$-normalized with a pseudo-count of 1. We found that by using $\mathbf{X}^*$ to derive GCS, as opposed to the original matrix $\mathbf{X}$, we could obtain a larger gain in performance over gene counts alone ($P$ = 0.006 vs. 0.04, respectively; paired two-sided $t$-test; **Fig. S8B**). This finding prompted us to apply this transformation as a pre-processing step for GCS and CytoTRACE throughout this work.

*Smoothing covariant gene expression across transcriptionally similar cells.*
Although GCS significantly improved the association with single-cell differentiation status (**Figs. S1B and S8A, B**), it was still observably noisy with considerable intra-phenotypic variance (e.g., **Fig. 3A, center**). Assuming transcriptionally similar cells occupy similar differentiation states, we applied a two-step smoothing procedure to improve upon GCS using nearest neighbor graphs.

First, we converted the normalized expression matrix $\mathbf{X}^*$ into a Markov process capturing local similarity between cells. In brief, $\mathbf{X}^*$ was filtered to a set of genes expressed in at least 5% of cells. The dispersion index was calculated for each gene by taking the ratio of its variance to its mean across all cells. The top 1,000 most dispersed genes were selected and used to compute a $k \times k$ similarity matrix, $\mathbf{D}$, where $\mathbf{D}_{ij}$ is the Pearson product-moment correlation coefficient between cells $i$ and $j$. The similarity matrix was then converted to a Markov matrix, $\mathbf{A}$, by setting all diagonal elements and correlations less than the null similarity coefficient $\partial$ to 0 and normalizing the sum of each row to 1. The null similarity coefficient $\partial$ was estimated by bootstrapping 10,000 random Pearson product-moment correlation coefficients between cells $i$ and $j$, where $i \neq j$. In practice, the null coefficient computed by bootstrapping was nearly identical to the mean of $\mathbf{D}$ after setting diagonal entries to zero. We therefore employed the latter approach to estimate $\partial$ (by default) to improve running time.

Next, we applied a two-step smoothing procedure to iteratively refine our estimate of the GCS vector (denoted $\overrightarrow{\mathrm{GCS}}$). First, we applied non-negative least squares regression (NNLS; R package: nnls v1.4) to solve the following optimization problem:

$$\arg\min_{\vec{\mathrm{x}} \geq 0} \parallel \mathbf{A}\vec{\mathrm{x}} - \overrightarrow{\mathrm{GCS}} \parallel_2$$

where $\vec{\mathrm{x}} \geq 0$ and $\overrightarrow{\mathrm{GCS}_\mathrm{R}} = \mathbf{A}\vec{\mathrm{x}}$. Because $\mathbf{A}$ is often collinear due to the presence of transcriptionally similar cells, the solution to NNLS, denoted by vector $\vec{\mathrm{x}}$, is generally sparse without requiring regularization or parameterization (unlike Lasso, for example[94]). As a result, the use of NNLS allows $\overrightarrow{\mathrm{GCS}}$ to be explained as a function of distinct transcriptional neighborhoods in $\mathbf{A}$ (denoted $\overrightarrow{\mathrm{GCS}_\mathrm{R}}$), resulting in a smoother representation and improved performance over $\overrightarrow{\mathrm{GCS}}$ (**Fig. S8B**). To further refine $\overrightarrow{\mathrm{GCS}_\mathrm{R}}$, we simulated a diffusion process, which was applied for 10,000 iterations (t = {1,…,10,000}) or until convergence ($\mathrm{mean}(|\vec{\mathrm{d}}_{\mathrm{t}+1} - \vec{\mathrm{d}}_\mathrm{t}|) \leq 1 \times 10^{-6}$):

$$\vec{\mathrm{d}}_{\mathrm{t}+1} = \alpha \cdot \mathbf{A}\vec{\mathrm{d}}_\mathrm{t} + (1 - \alpha) \cdot \vec{\mathrm{d}}_0$$

where $\vec{d}_0 = \overrightarrow{GCS_R}$ and $\alpha = 0.9$ (**Fig. S8D**). Unlike NNLS, which identifies nonnegative coefficients that best explain $\overrightarrow{GCS}$ as a function of $\mathbf{A}$, the diffusion process iteratively adjusts $\overrightarrow{GCS_R}$ based on the probability structure of the Markov process defined in $\mathbf{A}$. We found that combining the two approaches yielded better performance than either approach alone (**Fig. S8B**). As a final step, we converted the output of diffusion (i.e., vector $\vec{d}_{t+1}$) to rank-space, yielding $\overrightarrow{cytotrace}$, a vector of length *k* containing the predicted developmental ordering of every cell in $\mathbf{X}$.

13. Analysis of differentiation-associated genes

For the analysis of differentiation-associated genes in **Figure 3E**, we used the following datasets: 'Bone marrow (Smart-seq2)', 'Pancreatic alpha cell (C1)', 'Intestine (Drop-seq)', and 'Peripheral glia (Smart-seq2)' (see *'Data availability'*). All 42 gold standard datasets in this work were analyzed in **Figure S11**. Gene sets were defined as the 100 genes most enriched in the least differentiated ($rank_{min}$) or most differentiated ($rank_{max}$) cells by calculating the fold change of the average $\log_2$ expression of the former versus the remaining cells, and vice versa. All genes in each dataset were then rank-ordered by their Pearson correlation with CytoTRACE, and the Benjamini-Hochberg-adjusted *P* value for the enrichment of each gene set within the ranked transcriptome was calculated using the *clusterProfiler*[68] (v3.10.0) package in R with 'nPerm' set to 100,000.

14. 3D visualization of differentiation trajectories with CytoTRACE

CytoTRACE can usefully complement data visualization techniques, such as t-Distributed Stochastic Neighbor Embedding (tSNE)[95], Uniform Manifold Approximation and Projection (UMAP)[96], and force-directed layout algorithms[97], allowing visualization of predicted differentiation hierarchies. For example, to visualize whole zebrafish development and mouse bone marrow differentiation, CytoTRACE was added as a third axis to a 2D force-directed layout of each dataset (**Fig. 4A; Fig. S13**). Force-directed layouts were generated using the ForceAtlas2 [97] graphing algorithm, implemented as part of *scanpy*[98] v1.3.7 (Python). To visualize discontinuous cell states, such as during gastrulation and mesoderm differentiation (**Fig. S10A, C**), CytoTRACE was added as a third axis to a 2D layout of the first two principal components.

15. Combined root selection and branch detection with CytoTRACE and Monocle 2

States and branches were detected using the *monocle* v2.10.1 R package and following the instructions provided online (http://cole-trapnell-lab.github.io/monocle-release/docs/#constructing-single-cell-trajectories). Highly variable genes for trajectory inference were selected based on expression in at least 5% of cells, biological variability $\geq$ 0.5, and false discovery rate (FDR) $\leq$ 0.05 using the *scran* R package, as previously described[99]. Dimensional reduction was performed using the 'DDRTree' method and states were identified using the 'orderCells' function in the *monocle* v2.10.1 R package. The state with the highest mean pseudotime inferred by CytoTRACE was selected as the root, or the predicted origin of the differentiation process. This state was then indicated in the function 'orderCells' to generate a CytoTRACE-rooted pseudotime. Lineage-specific regulatory factors and marker genes were identified using the CytoTRACE-predicted pseudotimes and the Branched Expression Analysis Modeling (BEAM) platform in Monocle 2[34].

16. RNA velocity

RNA velocity[15] was run on two datasets with discontinuous cell states, 'Mesoderm (C1)' and 'hESC *in vitro* (C1)' (see *'Data availability'*), using *velocyto.py* v0.17.13 (Python) and *velocyto.R* v0.6 (R package) as per instructions provided online (velocyto.org). Raw reads were downloaded from the Sequence Read Archive (NCBI) and trimmed for base call quality (PHRED score $\geq$ 21) and adapter sequences using *skewer*[100] v0.2.2. Trimmed reads were then aligned to the hg38 reference genome (GRCh38.p12, version GCA_000001405.27) using *STAR*[101] v2.5.4b, generating sorted bam files. Loom files containing spliced, unspliced, and spanning reads were generated from the bam files using *velocyto.py* v0.17.13 (Python). Known phenotype labels from the original studies[32,39] were provided to enable default gene filtering by *velocyto.py*. Gene relative velocities were generated in *velocyto.R* and velocity fields were visualized using the 'pca.velocity.plot' function using recommended settings in the 'Chromaffin differentiation analysis tutorial' (http://pklab.med.harvard.edu/velocyto/notebooks/R/chromaffin2.nb.html).

17. Human subjects

All clinical specimens in this study were collected with informed consent for research use and were approved by the Stanford University and City of Hope Institutional Review Boards in accordance with the Declaration of Helsinki. These specimens, consisting of 6 tumor biopsies from patients with luminal-like breast cancer and 2 tumor biopsies from patients with basal-like breast cancer, were obtained from the primary site during surgical resection of breast tumors at Stanford Hospital and City of Hope National Medical Center. Paired adjacent normal tissues were also acquired for 7 of 8 patients. All samples were immediately dissociated for flow cytometry and sorting ('*Tissue dissociation*' below). In addition, tumor cells from two patients, COH69 and SU58, were lineage-depleted and transplanted into NOD *scid* gamma (NSG) mice to form xenografts, as described below.

18. Tissue dissociation

Fresh primary and xenografted tumors were mechanically dissociated into < 1-2 $mm^3$ pieces with a razor blade and then digested at 37 °C with 1500 U collagenase and 500 U hyaluronidase in Advanced DMEM/F12 (Thermo Fisher Scientific), 2 mM Glutamax (Invitrogen), and an antibiotic/antimycotic mix containing 120 µg/ml penicillin, 100 µg/ml streptomycin, and 0.25 µg/ml amphotericin-B (PSA) for 4-6 hrs with hourly pipetting for 5 min. After digestion, cells were treated with ACK lysis buffer to deplete red blood cells and then incubated with 10 U dispase to further dissociate tissue into single cells and 1000 U DNase I to prevent cell clumping. Cells were filtered through a 70 µm nylon mesh and washed with staining buffer containing 2% fetal bovine serum (FBS) and PSA in Hank's Balanced Salt Solution (HBBS).

19. Flow cytometry

Single cell suspensions of fresh breast tissue were stained with fluorescent antibodies to prepare for FACS. To reduce nonspecific antibody binding, single cells were blocked with 10 µg/mL rat IgG (Sigma-Aldrich) on ice for 10 minutes. Cells were then stained, in the dark, on ice for 30 minutes. FACS was performed with a 130 µm nozzle on a BD FACSAria II with BD FACSDiva software. For all experiments, side scatter and forward scatter profiles (area and width) were used to eliminate debris and cell doublets. Dead cells were eliminated by excluding 4',6-diamidino-2-phenylindole (DAPI) positive cells (Molecular Probes). For primary human breast samples, breast epithelial cells were enriched by negative gating of lineage cells expressing CD45, CD31, CD3, CD16, or CD64. Human breast basal cells were then gated as $CD49f^{high}EPCAM^{med\text{-}low}$, luminal progenitors as $CD49f^{high}EPCAM^{high}$, and mature luminal cells as $CD49f^{low}EPCAM^{high}$ (**Fig. S15A**).

For xenografted cells, human epithelial cells were enriched by negative gating of cells expressing H-2Kd, the murine MHC Class I molecule.

20. Single-cell expression profiling of human breast tumors

Single human breast cells were sorted into 96-well plates of lysis buffer as described above ('*Flow* cytometry') and cDNA libraries were constructed using a modified Smart-seq2 protocol and machine-automated pipeline by the Stanford Functional Genomic Facility (SFGF), as described previously[102,103]. Notably, 27 cycles of PCR amplification were required to obtain cDNA amounts within range of the recommended concentrations (0.05 – 0.32 ng/µL) for downstream tagmentation. Wells with over-amplified cDNA were diluted. Libraries were prepared using the Nextera XT DNA Library Prep (Illumina) kit and sequenced on a NextSeq 500 (Illumina) to obtain 2x75 bp paired-end reads.

21. Transcript quantification and cluster analysis of breast tumors

A total of 3,234 breast cells were sequenced from tumor and adjacent normal breast tissue, as described above. Raw FASTQ reads were aligned to GENCODE v25 reference transcripts (GRCh38.p7) with Salmon[84] v0.7.2 using the --seq-bias flag and otherwise default parameters. Salmon results were merged into a single gene-level TPM matrix using the R package, *tximport*[104] v1.4.0. Low quality cells, identified as cells expressing <1,000 detectable genes, were excluded from further analysis. Using Seurat[105], clusters were identified by running 'FindClusters' on the first 10 principal components of the data with the resolution parameter set to 0.6. Cell labels for known epithelial subpopulations were assigned according to the expression of previously described marker genes (*KRT14*$^{high}$, *MYLK*$^{high}$ = basal, *MMP7*$^{high}$, *RARRES1*$^{high}$ = luminal progenitor, *MUC1*$^{high}$, *AREG*$^{high}$ = mature luminal)[106,107]. Clusters with similar expression patterns of epithelial marker genes were merged and clusters lacking expression of expected markers were eliminated. These steps resulted in three remaining clusters that distinguished basal (tumor: $n$ = 294, normal: $n$ = 366), luminal progenitor (tumor: $n$ = 213, normal: $n$ = 319), and mature luminal cells (tumor: $n$ = 354, normal: $n$ = 356) (**Fig. S15B**). The final expression matrix was filtered to focus on protein-coding genes approved by the HUGO Gene Nomenclature Committee ($n$ = 19,031 genes).

22. Clonogenicity index

Related to **Figure 5B**, previously normalized gene expression microarrays (Illumina HumanHT-12 V4.0 beadchip) of sorted human epithelial subpopulations from normal breast tissues were downloaded (GSE35399)[54], and genes mapping to >1 probe were collapsed at the gene level to retain the probe with highest mean expression across all samples. Following quantile normalization, we defined a 'clonogenicity index' as the $\log_2$ fold change of each gene between ALDH+ LPs (more clonogenic) and ALDH- or ERBB3- LPs (less clonogenic).

23. RNAi dropout viability screen

Related to **Figure 5C**, genes that decreased human breast tumorigenesis ($n$ = 43) were chosen from a list of 46 hit candidates from an RNAi dropout viability screen[57]. *ASCC3L1, C15ORF15,* and *POLA* were excluded due to absence of expression in malignant LPs in our scRNA-seq data. For **Figure S16**, genes that did not decrease human breast tumorigenesis ($n$ = 50) were chosen from a list of 64 genes with the lowest shRNA ratios, or percent change from control shRNA. 14 genes were excluded due to absence of expression in malignant LPs in our scRNA-seq data. The normalized enrichment score (NES) and $Q$ value for the enrichment of each gene set within the

CytoTRACE-ranked gene list of malignant LPs was calculated using the *clusterProfiler*[68] v3.10.0 package in R.

24. Lentivirus production

Lentivirus was produced by transfecting 293T cells with Packaging Plasmid Mix (Cellecta) and subcloned pRSI12 shRNA expression plasmids using Lipofectamine 2000 (Thermo Fisher Scientific) per the manufacturer's instructions. Supernatants were collected at 48 hrs and 72 hrs, filtered with a 0.45 µm filter, and precipitated with Lentivirus Precipitation Solution (Alstem, LLC) per the manufacturer's instructions. Lentivirus was resuspended in 1/100 of the original volume. Viral titers were determined by flow cytometry analyses of RFP+ 293T cells infected with serial dilutions of concentrated virus.

25. Xenograft tumor cell transduction and engraftment

Dissociated single cells from xenografts were stained with biotin-conjugated anti-H-2Kd antibody and then anti-biotin microbeads (Miltenyl BioTec) to deplete mouse cells using autoMACS (Miltenyl BioTec). Tumor cells were transduced with pRSI12 (empty vector), shGULP1-1 (5'-TGCATACACCTGAAGCTTTAT-3'), and shGULP1-2 (5'-TCCTAAAGTGGAGTTGCAAAT-3'), at a MOI=25 for knockdown experiments. MOI was calculated based on infection in 293T cells using the formula Titre (TU/ml) = (N*P)/(V*D) where N=number of cells, P=Percentage infection (between 10-20%), V=volume of virus, D=Dilution fold. The transduced cells were washed and resuspended in staining media containing 50% matrigel and injected into the fourth abdominal fat pad by subcutaneous injection at the base of the nipple of female NSG mice (20,000 cells/mouse). Mice were monitored every week for tumor growth and measured using calipers to plot tumor volume V=L*W*H. The experiment was terminated when tumors from either control or knockdown reached 1500-2000 $mm^3$ in size.

26. Cell viability assay

Cell viability was assessed with MDA-MB-231, a human metastatic breast cancer cell line, after transduction with lentivirus containing an empty vector, *shGULP1-1*, or *shGULP1-2* at a seeding density of 5000 cells. Cell viability was measured at Day 0, 2, 4, and 7 by incubating cells with WST-1 Cell Proliferation Reagent (Roche) at 1:10 (v/v) final dilution at 37 $^{o}$C and 5% $CO_2$ for 1-4 hrs and analyzing absorbance at 450 nm on a SpectraMax M3 Bioanalyzer (Molecular Devices). To correct for the effect of media, background absorbance was measured in a blank well. Thus, the background corrected signal for each sample was $A_{corrected} = Absorbance_{experimental} – Absorbance_{background}$.

27. Statistical analysis

We calculated gene counts as the number of genes with detectable expression (>0 reads or UMIs). The concordance between known and predicted differentiation states was determined by an unweighted or weighted Spearman correlation (*rho*), depending on whether features were mean-aggregated by phenotype (unweighted) or not (weighted). Other linear relationships were modeled by linear regression ($R^2$), and a $t$ test was used to assess whether the result was significantly nonzero. When data were normally distributed, group comparisons were determined using a $t$ test with unequal variance or a paired $t$ test, as appropriate; otherwise, a Wilcoxon test was applied. Results with $P < 0.05$ were considered significant. Data analyses were performed with R, Prism v7 (GraphPad Software, Inc.), and FlowJo v10 (FlowJo, LLC). The investigators were not blinded to allocation during experiments and outcome assessment. No sample-size estimates were performed to ensure adequate power to detect a pre-specified effect size.

28. Software implementation and website

CytoTrace is implemented in R and is available as a standalone software tool or as a web application built using R Shiny and hosted at https://cytotrace.stanford.edu. The website runs on an Apache server on a virtual machine, and allows users to (1) perform custom CytoTRACE analyses using uploaded scRNA-seq expression matrices and phenotypic labels (if available), (2) view pre-computed CytoTRACE results from 42 benchmarking datasets, and (3) generate publication quality figures. The web interface features an interactive 3D scatter plot produced with Plotly that arranges single cell transcriptomes by a t-SNE projection on two axes and CytoTRACE on a third axis. By default, cells are colored by their CytoTRACE value, however options are provided to color the cells by (1) the expression of user-specified genes of interest and (2) cellular phenotype (if available). The website also generates boxplots of each user-defined phenotype ordered by CytoTRACE values and a plot showing the top ten genes predicted to be associated with the least differentiated and most differentiated cells. All plots and underlying data tables are downloadable. Finally, the web application provides (1) a vignette to illustrate the use of CytoTRACE, (2) answers to common questions, and (3) links to download the CytoTRACE source code and benchmark datasets.

29. Code availability

CytoTRACE is freely available for non-profit academic use at https://cytotrace.stanford.edu.

30. Data availability

The following datasets are available from the Gene Expression Omnibus (GEO): GSE59114 ('Aging HSCs (Smart-seq2)'[69]), GSE74767 ('Blastocyst phenotypes (SC3-seq)' and 'Blastocyst timepoints (SC3-seq)'[73]), GSE90860 ('Cortical interneurons (C1)'[108]), GSE95753 ('Dentate gyrus phenotypes (10x)' and 'Dentate gyrus timepoints (10x)'[33]), GSE67123 ('Embryonic HSCs (Tang et al.)'[109]), GSE98451 ('Endometrium (CEL-seq)'[110]), GSE99933 ('Peripheral glia (Smart-seq2)'[111], GSE94641 ('Medial ganglionic eminence (C1)'[112]), GSE60783 ('Dendritic cells (C1)'[87]), GSE67602 ('Hair epidermis (C1)'[113]), GSE70245 ('HSPCs (C1)'[30]), GSE90047 ('Hepatoblast (Smart-seq2)'[114]), GSE75748 ('hESC *in vitro* (C1)'[39]), GSE52529 ('HSMM (C1)'[29]), GSE85066 ('Mesoderm (C1)' [32]), GSE93421 ('Peripheral blood (10x)'[75]), GSE36552 ('Pre-implant human embryo (Tang et al.)'[78]), GSE86146 '(Germ cells (Smart-seq2)'[115]), GSE98664 ('mESC *in vitro* (RamDA-seq)'[116]), GSE52583 ('Lung development (C1)' and 'AT2/AT1 lineage (C1)'[27]), GSE97391 ('Direct *in vitro* neuron (inDrop)' and 'Standard *in vitro* neuron (inDrop)'[117]), GSE76408 ('Lgr5-CreER intestine (CEL-seq)'[18]), GSE92332 ('Intestine (Smart-seq2)' and 'Intestine(Drop-seq)'[43]), GSE45719 ('Pre-implant mouse embryo (Tang et al.)'[86]), GSE69761 ('Lung fibroblast (C1)'[70]), GSE107122 ('Neural stem cells (Drop-seq)'[76]), GSE102066 ('*in vitro* NPCs (C1)'[26]), GSE75330 ('Oligodendrocyte phenotypes (C1)' and 'Oligodendrocyte timepoints (C1)'[118]), GSE87375 ('Pancreatic alpha cell (Smart-seq2)' and 'Pancreatic beta cell (Smart-seq2)'[119]), GSE103633 ('Whole planaria (Drop-seq)'[74]), GSE107910 ('Thymus (Drop-seq)'[77]), GSE106587 ('Early zebrafish (Drop-seq)'[44]), and GSE109774 ('Bone marrow (Smart-seq2)' and 'Bone marrow (10x)'[42]). The 'Skeletal stem cells (C1)' dataset is available upon request from

C.K.F. Chan and colleagues[120]. Single-cell RNA-seq expression data generated in this study are hosted at https://cytotrace.stanford.edu with a GEO accession code pending.

**A**

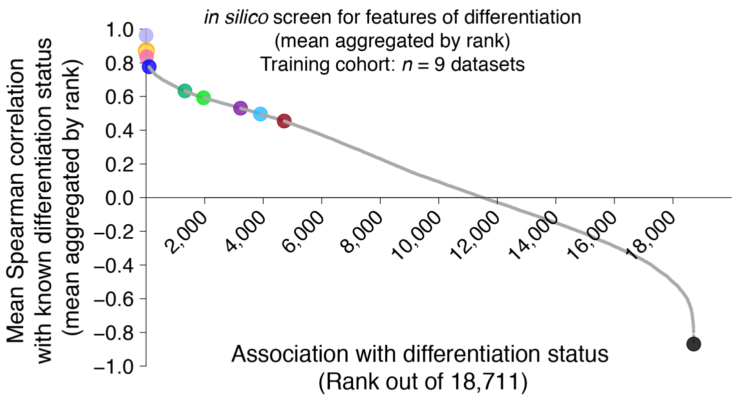


| Color | Features | Type | Mean ρ | Nominal Rank | Nominal Percentile | Absolute Rank | Absolute Percentile |
|---|---|---|---|---|---|---|---|
| | GO_SIGNAL_TRANSDUCTION_BY_P53_CLASS_MEDIATOR | Geneset | 0.96 | 1 | 99.99 | 1 | 99.99 |
| | GO_CHROMATIN_BINDING | Geneset | 0.87 | 7 | 99.96 | 7 | 99.96 |
| | GO_REGULATION_OF_CELL_CYCLE_G1_S_PHASE_TRANSITION | Geneset | 0.86 | 9 | 99.95 | 10 | 99.95 |
| | GO_REGULATION_OF_STEM_CELL_POPULATION_MAINTENANCE | Geneset | 0.84 | 15 | 99.92 | 17 | 99.91 |
| | GENE COUNTS | Biological | 0.78 | 104 | 99.44 | 112 | 99.40 |
| | MUELLER_PLURINET | Geneset | 0.63 | 1,328 | 92.90 | 1,471 | 92.14 |
| | SLICE entropy | Method | 0.59 | 1,964 | 89.50 | 2,216 | 88.16 |
| | SCENT entropy | Method | 0.53 | 3,227 | 82.75 | 3,750 | 79.96 |
| | mRNA stemness index (mRNAsi) | Method | 0.50 | 3,907 | 79.12 | 4,643 | 75.19 |
| | StemID | Method | 0.45 | 4,719 | 74.78 | 5,742 | 69.31 |
| | GO_DISRUPTION_OF_CELLS_OF_OTHER_ORGANISM | Geneset | -0.87 | 18,711 | 0.00 | 9 | 99.95 |

**B**

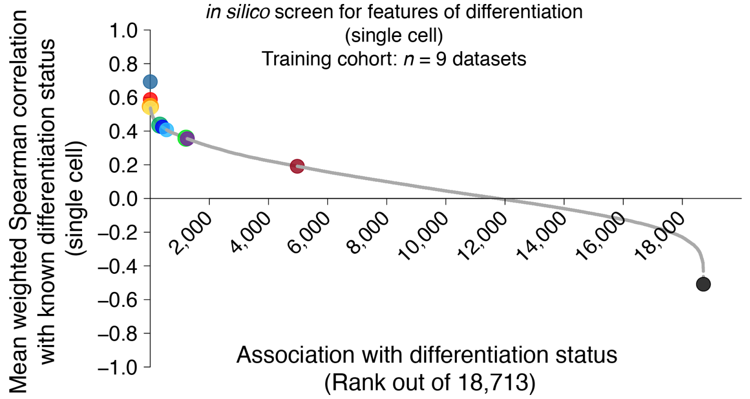


| Color | Features | Type | Mean ρ | Nominal Rank | Nominal Percentile | Absolute Rank | Absolute Percentile |
|---|---|---|---|---|---|---|---|
| | CytoTRACE | Method | 0.69 | 1 | 99.99 | 1 | 99.99 |
| | Gene counts signature (GCS) | Geneset | 0.59 | 2 | 99.99 | 2 | 99.99 |
| | GO_CHROMATIN_BINDING | Geneset | 0.55 | 3 | 99.98 | 3 | 99.98 |
| | GO_CELL_CYCLE | Geneset | 0.54 | 8 | 99.96 | 8 | 99.96 |
| | MUELLER_PLURINET | Geneset | 0.44 | 325 | 98.26 | 328 | 98.25 |
| | GENE COUNTS | Biological | 0.42 | 414 | 97.79 | 419 | 97.76 |
| | mRNA stemness index (mRNAsi) | Method | 0.41 | 551 | 97.06 | 560 | 97.01 |
| | SLICE | Method | 0.36 | 1,211 | 93.53 | 1,263 | 93.25 |
| | SCENT | Method | 0.35 | 1,254 | 93.30 | 1,313 | 92.98 |
| | StemID | Method | 0.19 | 4,975 | 73.41 | 6,236 | 66.68 |
| | HALLMARK_MYOGENESIS | Geneset | -0.51 | 18,713 | 0.00 | 39 | 99.79 |

**Figure S1. *In silico* screen for features of differentiation.** (**A**) *Top*: Same as in **Fig. 1C**. *Bottom*: Table summary of key features from the phenotype-level screen in the training cohort (*n* = 9; **Methods**). (**B**) Same as in **A** but evaluating performance at the single-cell level, which was calculated as the mean weighted (by number of cells per phenotype) Spearman correlation between the single cell values of each feature and known differentiation status across nine datasets.

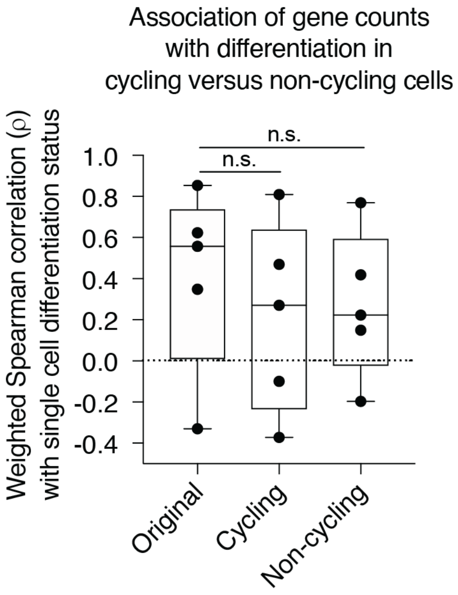


**Figure S2 Association between gene counts and differentiation in cycling and non-cycling cells.** Boxplots showing the weighted (by number of cells per phenotype) Spearman correlation between gene counts and single cell differentiation status (*y*-axis) in unstratified, cycling, and non-cycling single cells (*x*-axis). Only datasets in the training cohort with bimodal cell cycle gene expression and at least 10 cycling or non-cycling cells ($n = 5$) are shown. Statistical significance was assessed by a paired Wilcoxon signed-rank test. n.s. = not significant ($P > 0.05$). For further details, see **Methods**.

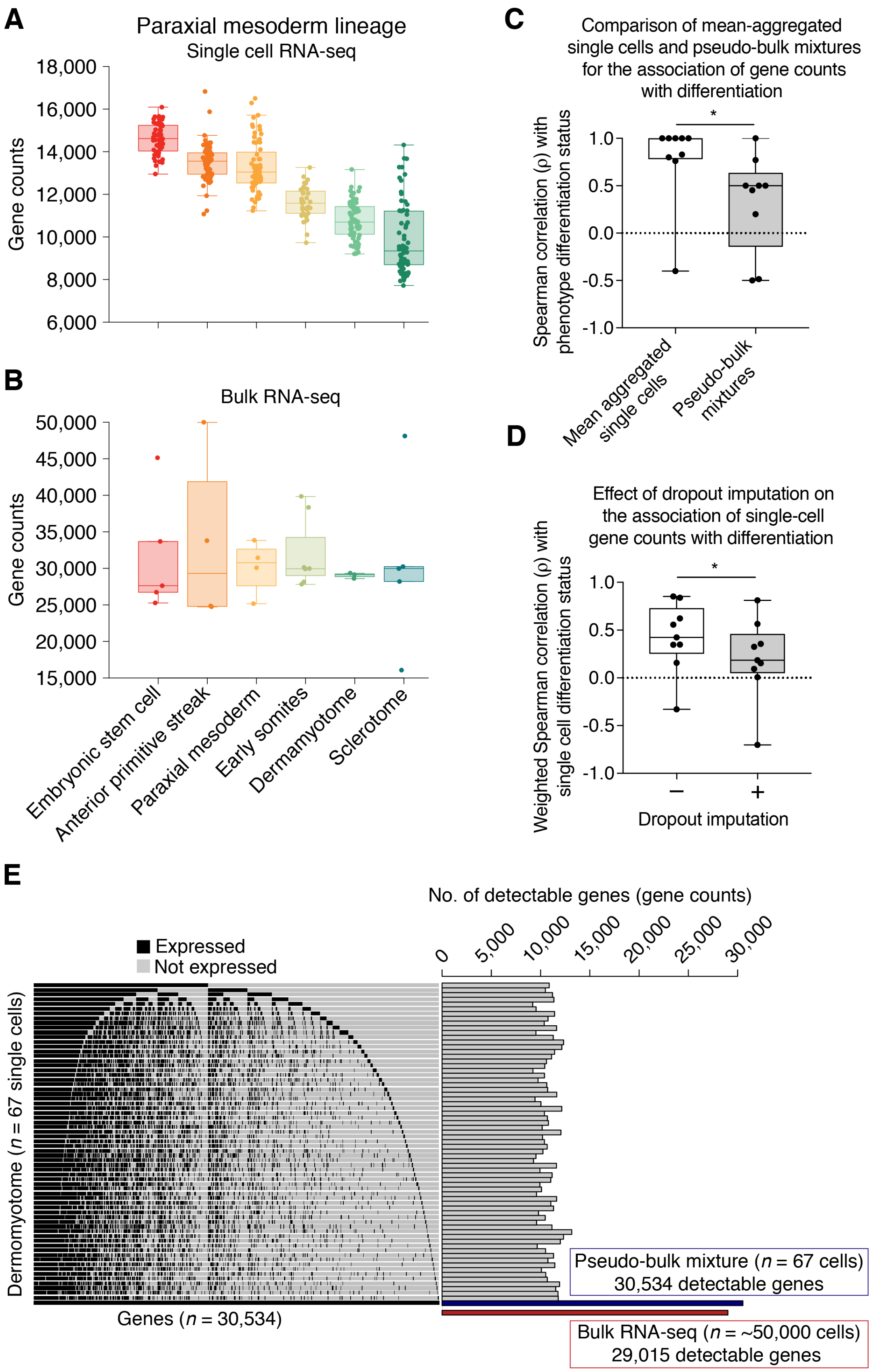
A
Paraxial mesoderm lineage
Single cell RNA-seq
Gene counts
18,000
16,000
14,000
12,000
10,000
8,000
6,000
B
Bulk RNA-seq
Gene counts
50,000
45,000
40,000
35,000
30,000
25,000
20,000
15,000
Embryonic stem cell
Anterior primitive streak
Paraxial mesoderm
Early somites
Dermamyotome
Sclerotome
C
Comparison of mean-aggregated single cells and pseudo-bulk mixtures for the association of gene counts with differentiation
Spearman correlation (ρ) with phenotype differentiation status
*
1.0
0.5
0.0
-0.5
-1.0
Mean aggregated single cells
Pseudo-bulk mixtures
D
Effect of dropout imputation on the association of single-cell gene counts with differentiation
Weighted Spearman correlation (ρ) with single cell differentiation status
*
1.0
0.5
0.0
-0.5
-1.0
−
+
Dropout imputation
E
No. of detectable genes (gene counts)
0
5,000
10,000
15,000
20,000
25,000
30,000
Expressed
Not expressed
Dermomyotome (n = 67 single cells)
Genes (n = 30,534)
Pseudo-bulk mixture (n = 67 cells)
30,534 detectable genes
Bulk RNA-seq (n = ~50,000 cells)
29,015 detectable genes

**Figure S3 Approaches to overcome sparsity in scRNA-seq data degrade the predictive performance of gene counts.** (**A-E**) Comparison of single-cell gene counts with gene counts from bulk RNA-seq, pooled single-cell, and dropout-imputed transcriptomes. (**A**) Boxplots showing the association of gene counts from single-cell transcriptomes (*y*-axis) with differentiation (*x*-axis) during paraxial mesoderm differentiation from the 'Mesoderm (C1)' dataset (**Methods**). (**B**) Boxplots showing the association of gene counts from bulk RNA-sequencing profiles (*y*-axis) with differentiation (*x*-axis) during paraxial mesoderm differentiation from the same study in **A**. (**C**) Boxplots comparing the association of mean single-cell gene counts per phenotype and gene counts of pooled single-cell transcriptomes per phenotype with differentiation. Spearman correlations between gene counts and known differentiation status for each dataset ($n$ = 9) are plotted as individual points. Statistical significance was assessed by a one-sided paired Wilcoxon signed-rank test; *$P < 0.05$. (**D**) Boxplots showing the effect of dropout imputation in single-cell transcriptomes on the association of single-cell gene counts with differentiation. Weighted (by number of cells per phenotype) Spearman correlations for each dataset ($n$ = 9) are plotted as individual points. Statistical significance was assessed by a one-sided paired Wilcoxon signed-rank test; *$P < 0.05$. (**E**) *Left*: Heatmap showing differential sampling of 30,534 genes (*x*-axis) by 67 single dermomyotome cells. Expressed (TPM >0) genes are in black, and non-expressed (TPM = 0) genes are in gray. The OncoPrint algorithm[88] was applied to show distinct patterns of transcriptome sampling. *Right:* Bar plots showing gene counts in single dermomyotome cells (gray), gene counts after pooling all 67 single cell transcriptomes (blue), and mean gene counts from 3 bulk RNA-seq transcriptomes (as shown in **B**) of ~50,000 purified dermomyotome cells each (red). For additional details, see **Methods**.

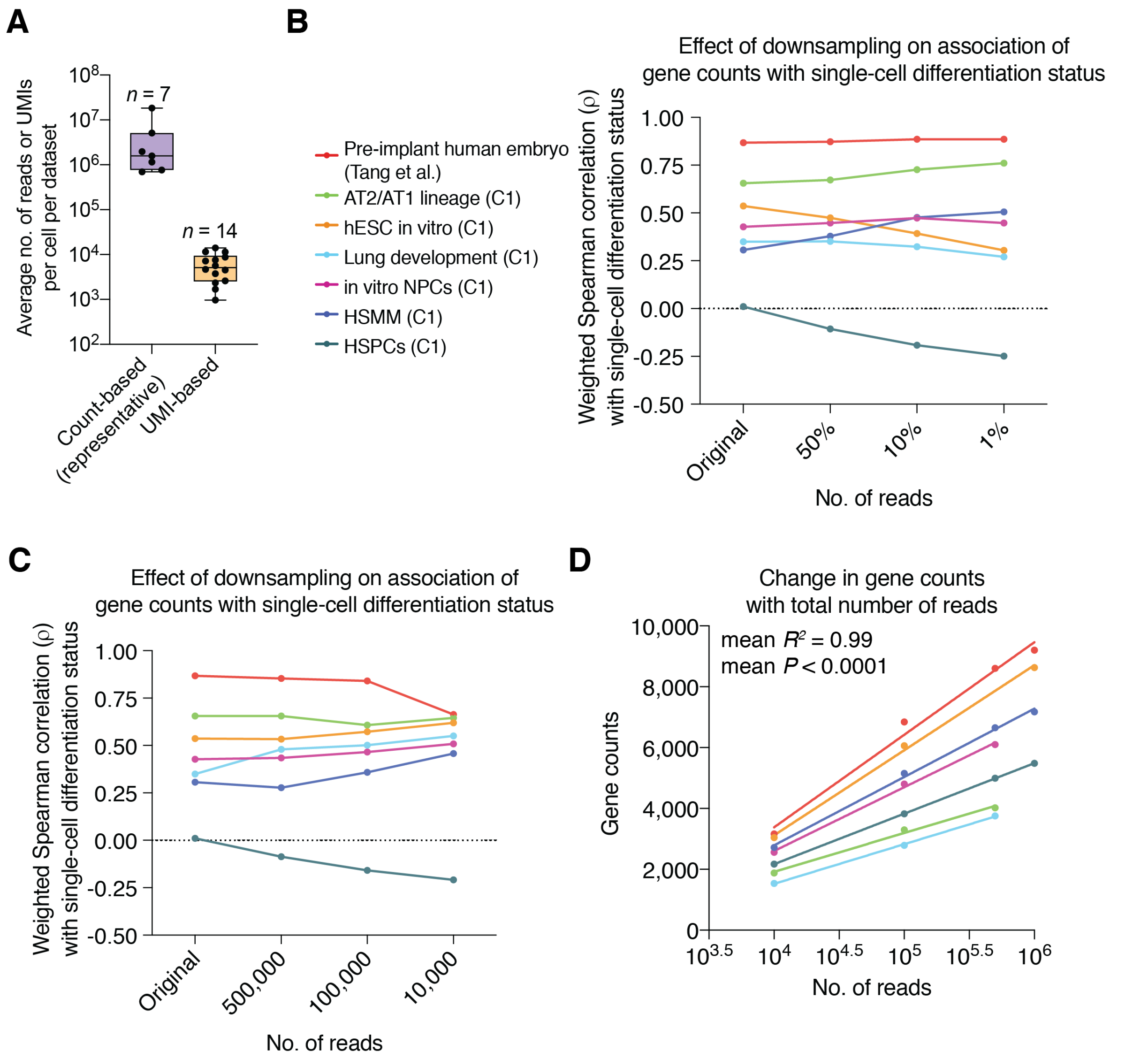


**Figure S4 Impact of the number of reads per cell on gene counts.** (**A**) Box plots comparing the average number of reads/cell or UMIs/cell for datasets analyzed in this study. Evaluable datasets related to the former are listed in panel B. (**B**) Line plot showing effect of reads down-sampling on the association of gene counts with single cell differentiation. Total number of reads per single cell were down-sampled to 50%, 10%, and 1% of original number of reads and the weighted (by number of cells per phenotype) Spearman correlation between gene counts and known single cell differentiation status was calculated for each dataset shown. (**C**) Line plot showing effect of reads down-sampling on the association of gene counts with single cell differentiation. Total number of reads per single cell were down-sampled to 500,000, 100,000, and 10,000 of original number of reads and the weighted (by number of cells per phenotype) Spearman correlation between gene counts and known single cell differentiation status was calculated for each dataset shown. (**D**) Linear regression lines showing the association between the logarithmic number of reads and mean gene counts per dataset using down-sampled transcriptomes. Mean $R^2$ of the regressions and mean $P$ values as determined by a $t$-test are indicated.

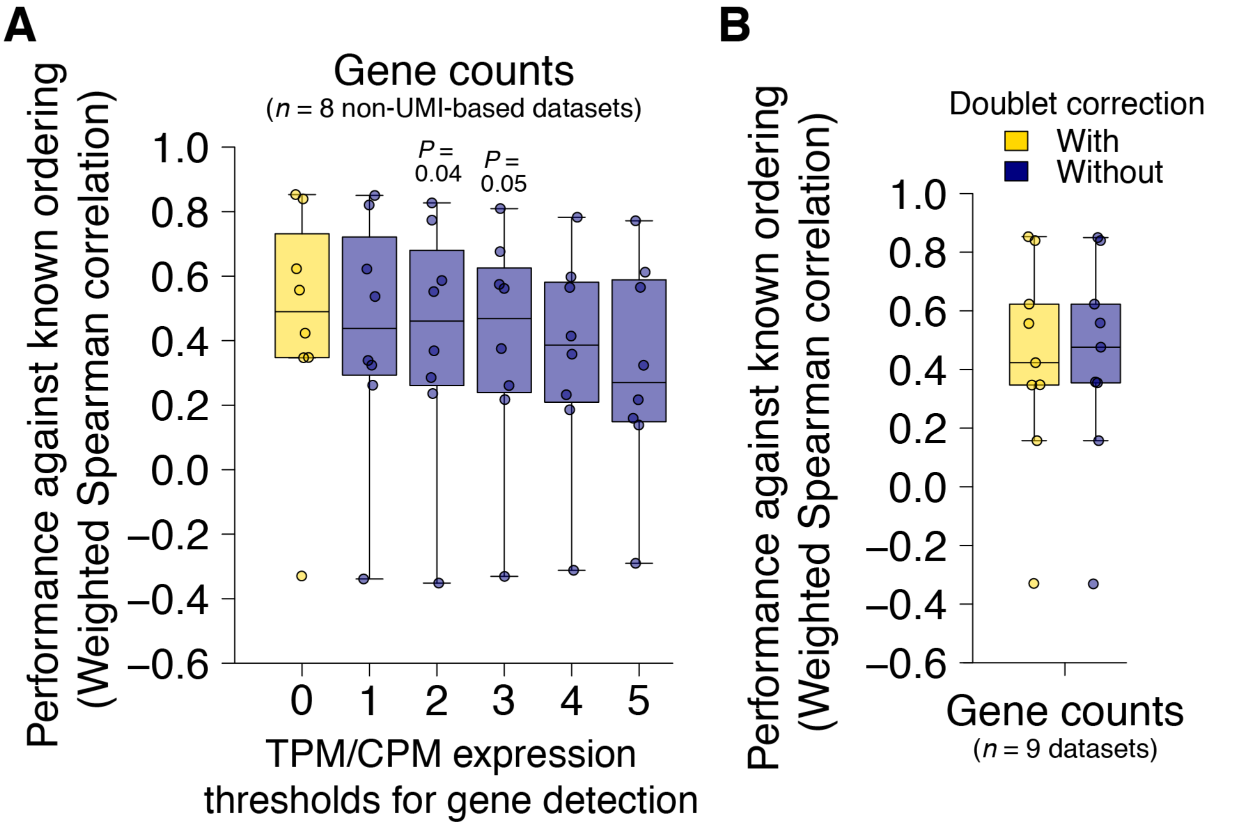


**Figure S5 Impact of different minimum expression thresholds and doublet removal on the association between gene counts and developmental potential.** (**A**) Boxplots showing the effect of increasing the minimum expression threshold for defining the number of genes per cell on the association between gene counts and developmental potential. The number of expressed genes per cell was mean-aggregated by phenotype prior to analysis and only non-UMI-based datasets in the training cohort (**Fig. 1A**) were considered ($n$ = 8). Statistical significance was assessed by a two-sided paired Wilcoxon signed-rank test against TPM/CPM>0 (highlighted in yellow). TPM, transcripts per million; CPM, counts per million. (**B**) Boxplots showing the effect of removing doublets on the association between gene counts and developmental potential. Doublets were detected in the training cohort ($n$ = 9) using the *Scrublet* v0.2 package in Python with default settings[121].

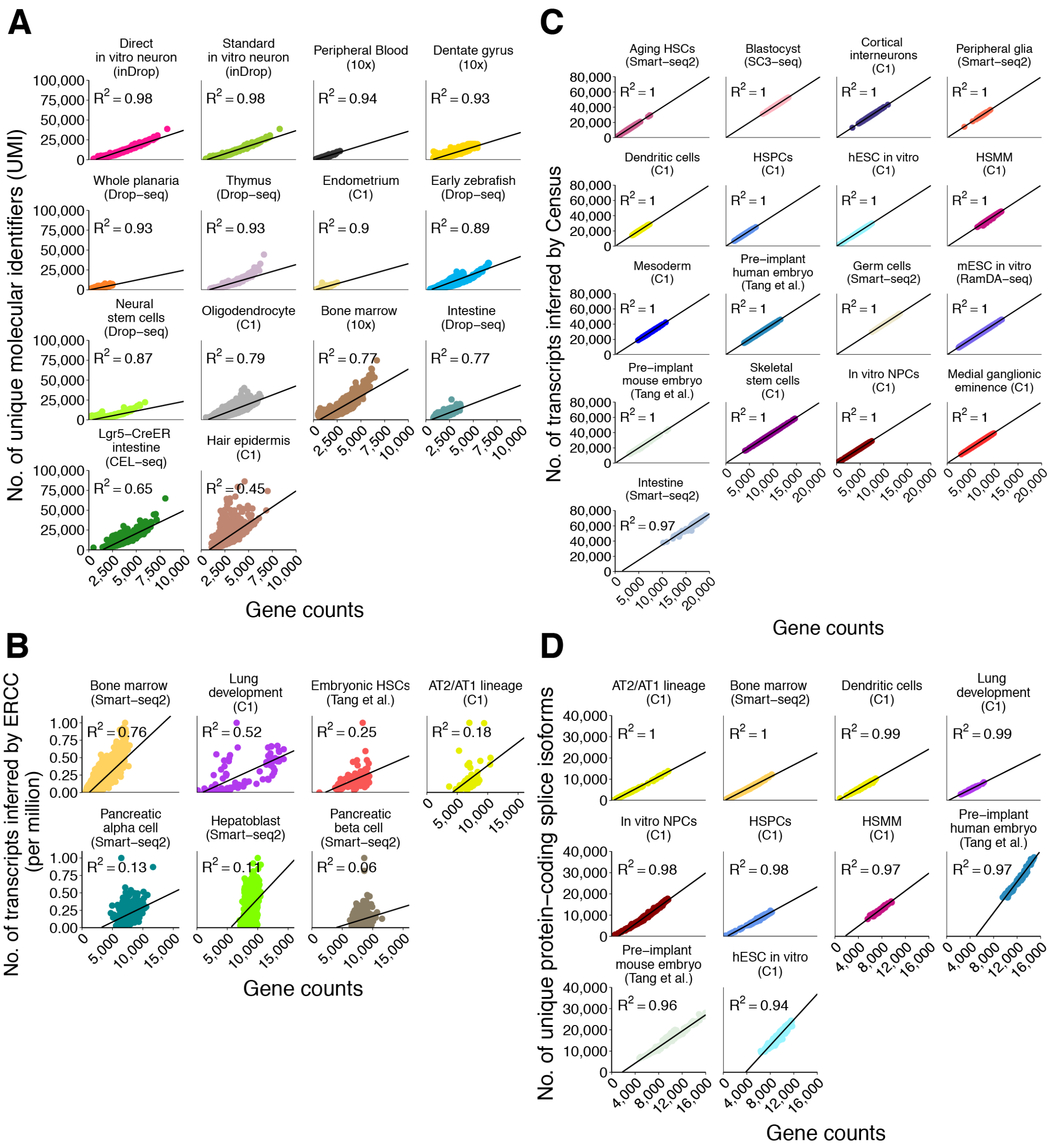


**Figure S6 Association between gene counts and RNA abundance.** (**A**) Scatterplots showing the association of gene counts with total number of unique molecular identifiers (UMIs) per single cell for datasets in this study where this information was available (*n* = 14). (**B**) Scatterplots showing the association of gene counts with the absolute number of detectable transcripts per cell. The latter was inferred by a linear regression model using 92 ERCC spike-ins for datasets in this study where this information was available (*n* = 7; *'Analysis of total RNA content and transcriptional diversity'* in **Methods**). Coefficient of determination, or $R^2$, is indicated for each dataset. (**C**) Scatterplots showing the association of gene counts with the absolute number of transcripts per cell. The latter was inferred by Census for all non-UMI datasets in this study that lack external ERCC standards (*n* = 17). (**D**) Scatterplots showing the association of gene counts with the number of unique protein-coding splice isoforms for all remapped datasets in this study (*n* = 10). Coefficient of determination, or $R^2$, is indicated for each dataset in **A-D**.

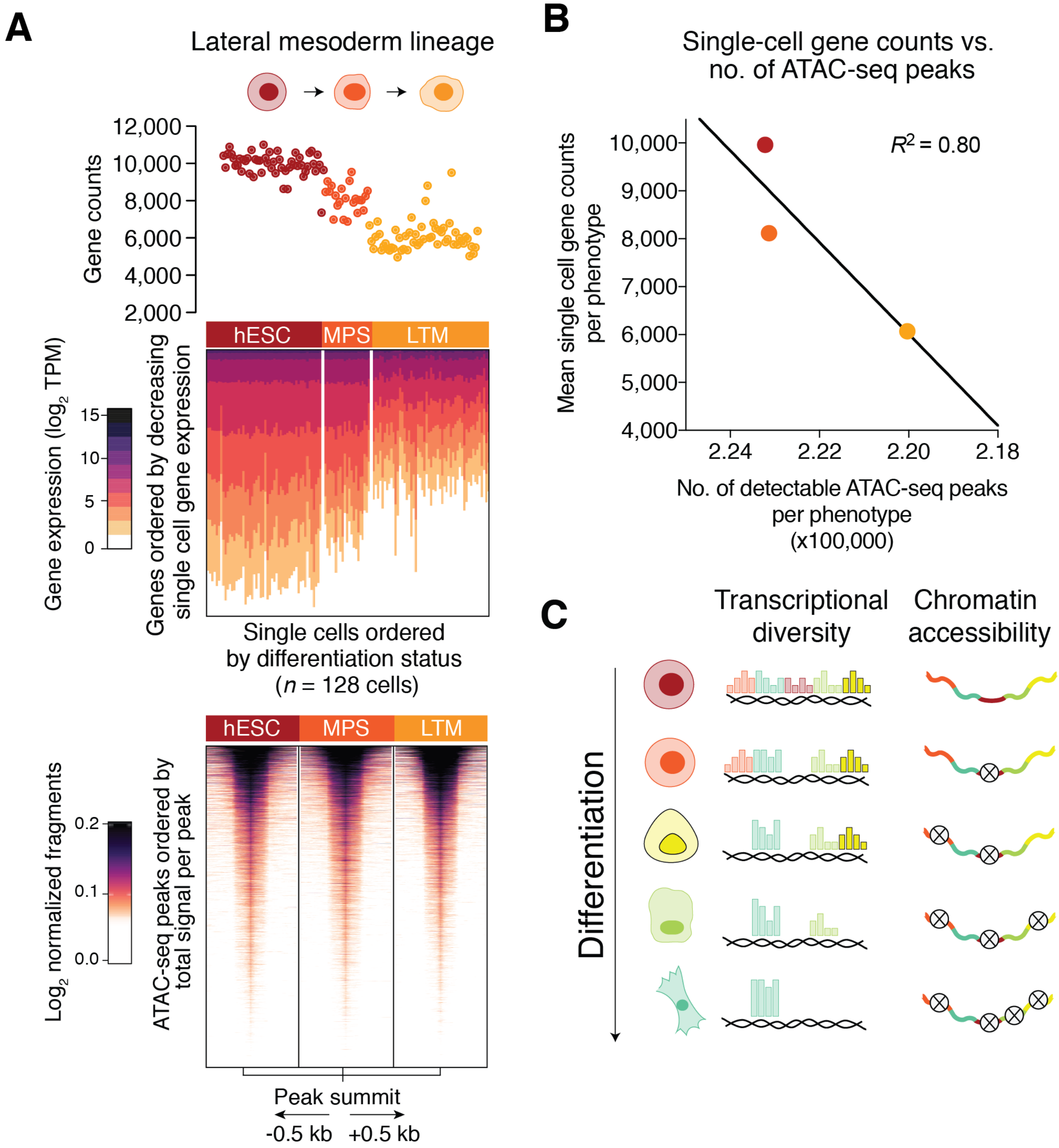


**Figure S7 Dynamics of gene counts, transcriptional diversity, and global chromatin accessibility during *in vitro* differentiation of hESCs into lateral mesoderm.** (**A, B**) Same as **Fig. 2B, C**, but showing the lateral mesoderm lineage from the 'Mesoderm (C1)' dataset (**Methods**). (**C**) Model of the decrease in transcriptional diversity and chromatin accessibility during cellular differentiation.

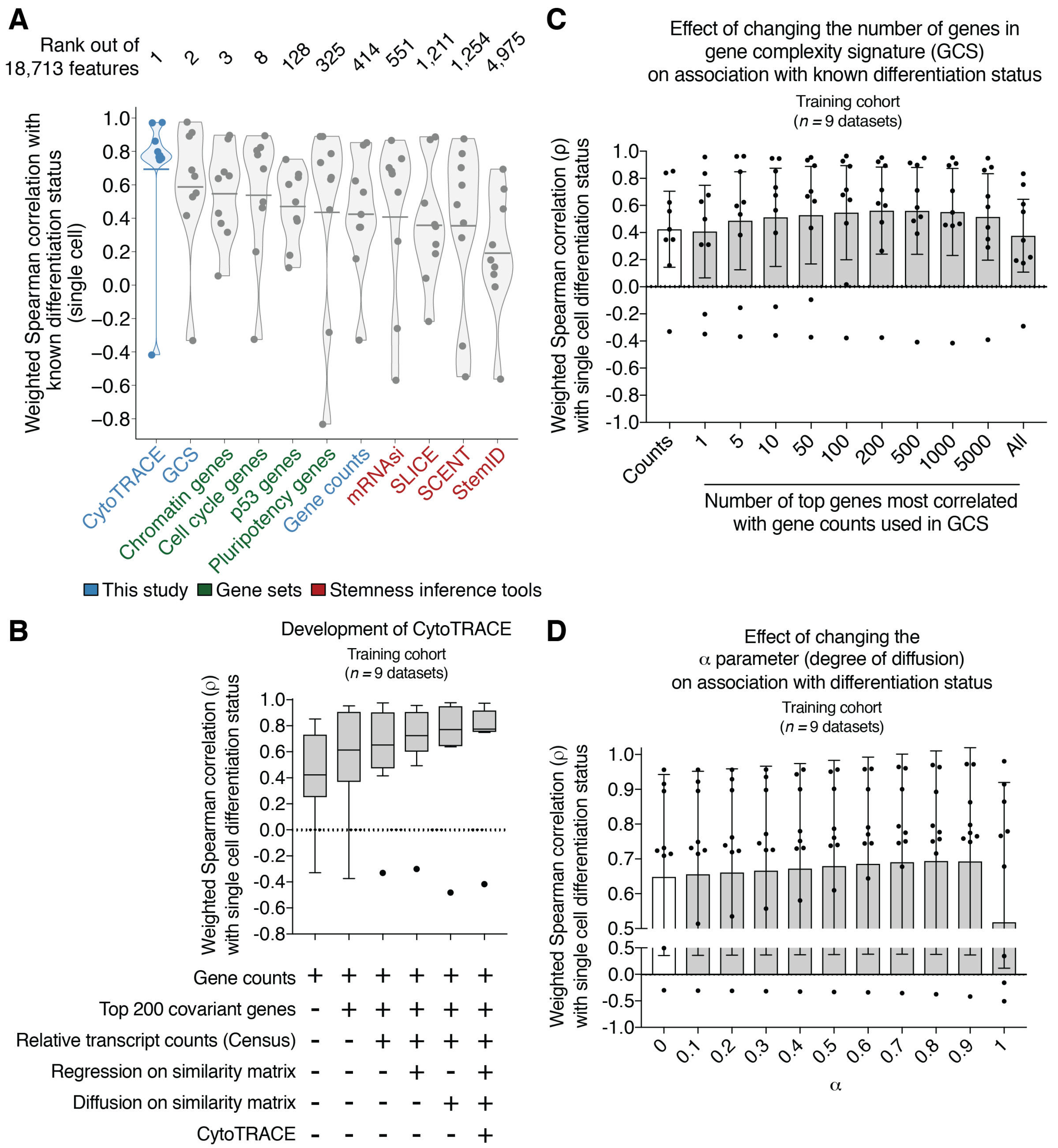


**Figure S8 Development and robustness of CytoTRACE.** (**A**) Single-cell level performance of CytoTRACE, gene counts signature (GCS), gene counts, entropy-based methods for stemness prediction, mRNA stemness index (mRNAsi), and the top differentiation-associated gene sets in the nine datasets from the training cohort. Data are presented as violin plots with horizontal bars indicating means. Features and computational strategies are ordered left to right by mean. (**B**) Tukey box plots showing the change in association with known differentiation status with sequential development of CytoTRACE from gene counts (**Methods**). (**C**) Bar plots showing the effect of the number of covariant genes used to calculate the gene counts signature (GCS) on association with known differentiation status (**Methods**). (**D**) Bar plots showing the effect of changing the α parameter, which affects the degree of diffusion applied to the input, on the association with known differentiation status. Data analyzed in **A-D** are from the training cohort (**Fig. 1A**).

**A**

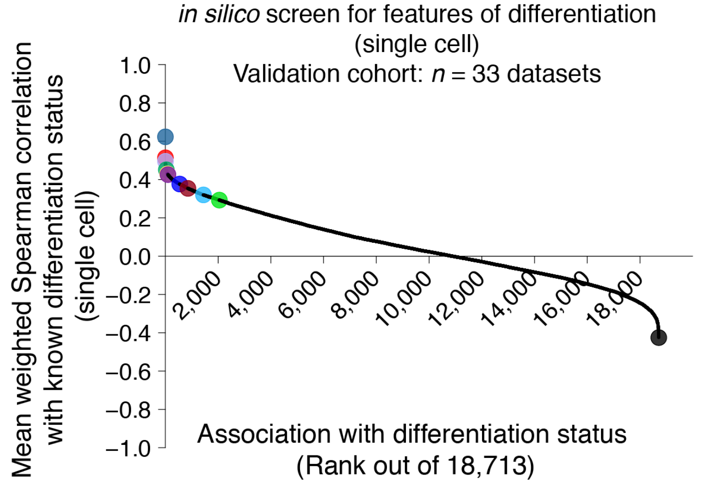


| Color | Features | Type | Mean ρ | Nominal | | Absolute | |
|---|---|---|---|---|---|---|---|
| | | | | Rank | Percentile | Rank | Percentile |
| | CytoTRACE | Method | 0.62 | 1 | 99.99 | 1 | 99.99 |
| | Gene counts signature (GCS) | Geneset | 0.51 | 2 | 99.99 | 2 | 99.99 |
| | GO_RNA_PROCESSING | Geneset | 0.50 | 3 | 99.98 | 3 | 99.98 |
| | MUELLER_PLURINET | Geneset | 0.45 | 35 | 99.81 | 35 | 99.81 |
| | GO_CELL_CYCLE | Geneset | 0.43 | 82 | 99.56 | 82 | 99.56 |
| | SCENT | Method | 0.43 | 98 | 99.48 | 98 | 99.48 |
| | GENE COUNTS | Biological | 0.38 | 544 | 97.09 | 560 | 97.01 |
| | StemID | Method | 0.35 | 854 | 95.44 | 885 | 95.27 |
| | mRNA stemness index (mRNAsi) | Method | 0.32 | 1,444 | 92.28 | 1,566 | 91.63 |
| | SLICE | Method | 0.29 | 2,047 | 89.06 | 2,305 | 87.68 |
| | chr21p11 | Geneset | -0.43 | 18,713 | 0.00 | 100 | 99.47 |

**B**

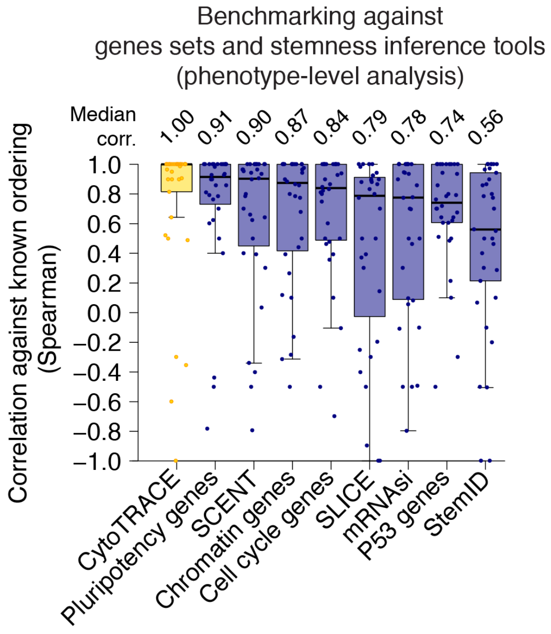


**Figure S9 Prediction of single-cell differentiation states in 33 validation datasets.** (**A**) Same as in **Figure 1C,** but evaluating performance at the single-cell level and in the validation cohort (*n* = 33 datasets). (**B**) Same as in **Figure 3D**, but comparing phenotype-level performance of CytoTRACE, top-performing gene sets, and stemness inference tools in the validation cohort. For further details, see **Methods**.

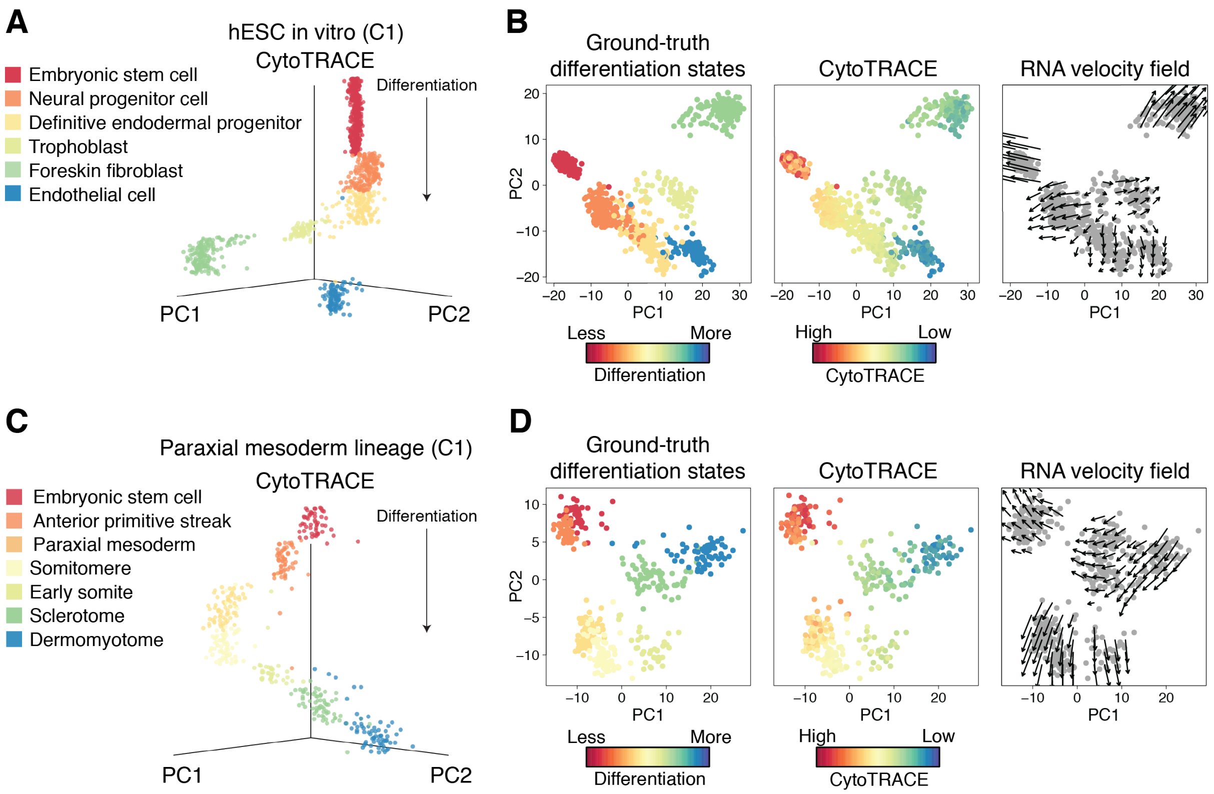


**Figure S10 Prediction of single-cell differentiation states without transitional cells.** (**A-D**) Potential application of CytoTRACE in datasets without transitional cell states. (**A**) 3D scatter plots of *in vitro* gastrulation process ('hESC *in vitro* (C1)' dataset; **Methods**), showing the predicted ordering of single cells by CytoTRACE in relation to the first two principal components, PC1 and PC2. Known differentiation is indicated by color. (**B**) Plots of PC1 and PC2 from **A**, showing, from left to right, ground-truth differentiation states (color), CytoTRACE (color), and the predicted future state of each cell determined by RNA velocity (**Methods**). (**C, D**) Same as in **A, B**, but with single cells from hESC differentiation through the paraxial mesoderm lineage ('Mesoderm (C1)'; **Methods**).

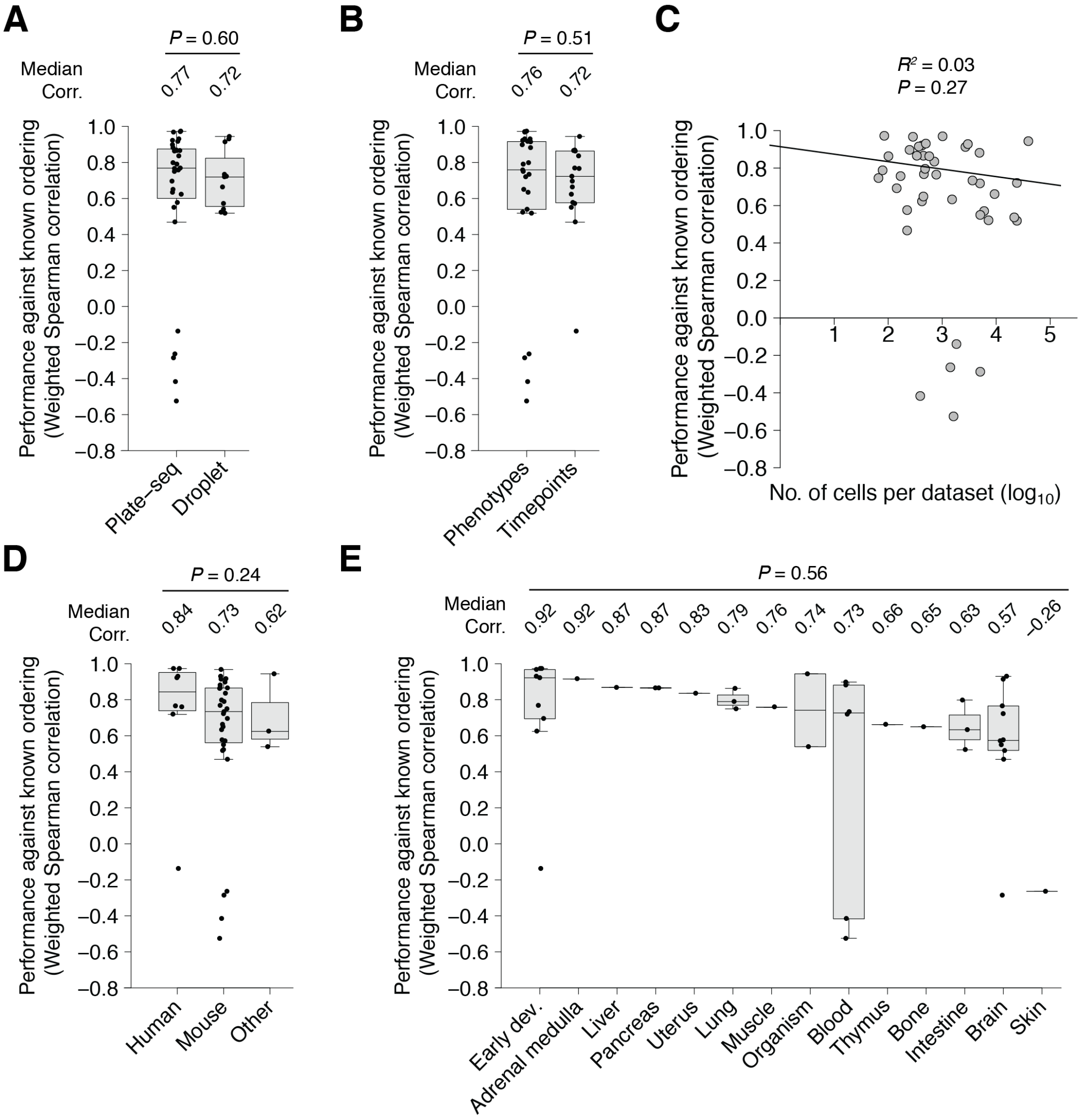


**Figure S11 Robustness of CytoTRACE to variation in dataset characteristics.** (**A-E**) Predictive performance of CytoTRACE across platforms, species, tissues, and experimental designs for all datasets in this study (*n* = 42; **Methods**). Performance was evaluated as the weighted (by number of cells per phenotype) Spearman correlations between CytoTRACE and known differentiation status. (**A**) Boxplots comparing the performance of CytoTRACE in scRNA-seq datasets prepared by plate-based versus droplet-based methods. Statistical significance was assessed by an unpaired Wilcoxon signed-rank test. (**B**) Boxplots comparing the performance of CytoTRACE in scRNA-seq datasets describing development over a time series versus differentiation of known phenotypes. Statistical significance was assessed by an unpaired Wilcoxon signed-rank test. (**C**) Scatter plot showing the association between the number of cells per dataset (*x*-axis) and CytoTRACE performance (*y*-axis). Statistical significance was assessed by a linear regression *t*-test. (**D**) Boxplots comparing the performance of CytoTRACE in scRNA-seq datasets from mouse, human, or other species (e.g. macaque, zebrafish, and planaria). Statistical significance was assessed by a Kruskal-Wallis rank sum test. (**E**) Boxplots comparing the performance of CytoTRACE in scRNA-seq datasets from 14 different tissues. Statistical significance was assessed by a Kruskal-Wallis rank sum test.

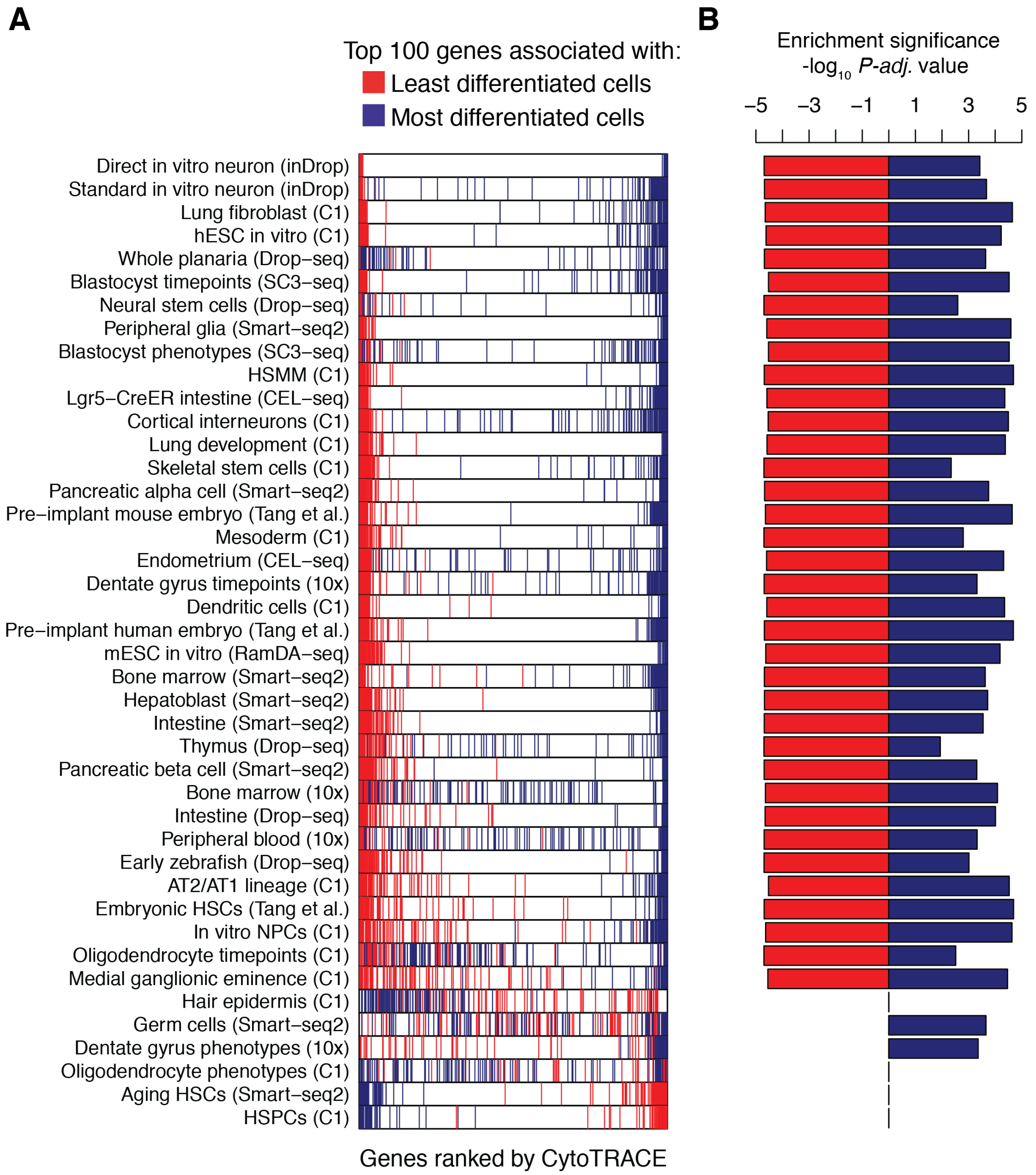


**Figure S12 Evaluation of CytoTRACE for prioritizing developmental marker genes.** (**A**) Plots showing the enrichment of key stemness-associated (red) and differentiation-associated (blue) genes by CytoTRACE in the full cohort (*n* = 42 datasets, **Methods**). Top 100 genes were defined as the 100 genes most enriched in the least differentiated or most differentiated cells by $\log_2$-fold change. (**B**) Statistical significance of the results on the left, as calculated by gene set enrichment analysis using the *clusterProfiler* package in R. Values are represented as $-\log_{10}$ Benjamini-Horchberg-adjusted *P* value (*P-adj*).

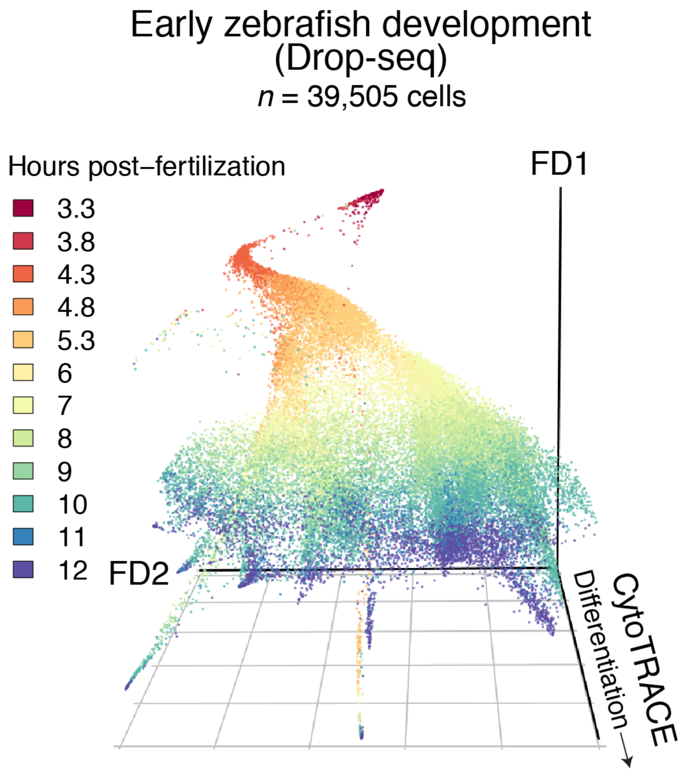


**Figure S13 Reconstruction of early zebrafish development.** 3D scatter plot of early zebrafish development (Drop-seq; **Methods**), showing single-cell CytoTRACE predictions versus a force-directed layout (FD1 and FD2). Post-fertilization time points are indicated by color.

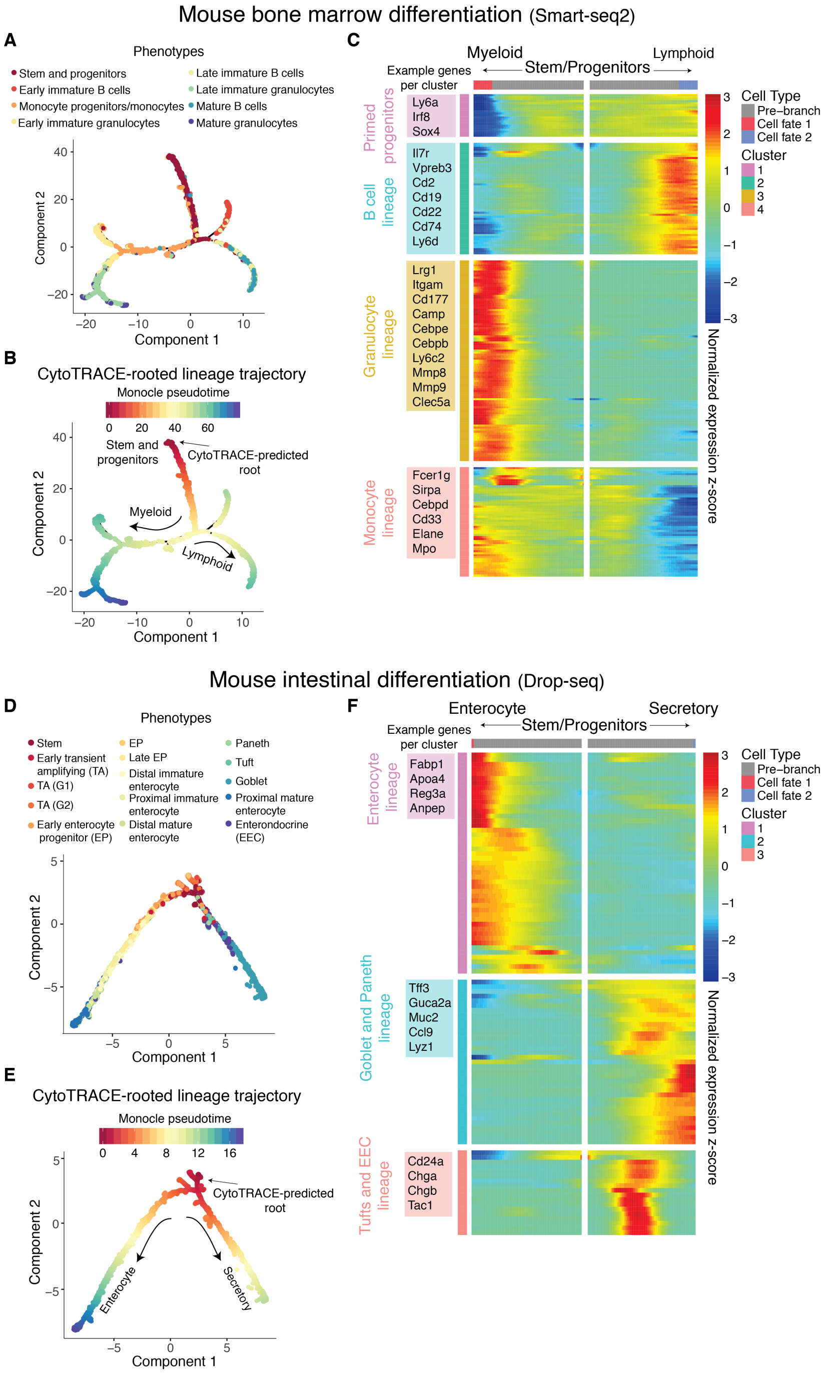


**Figure S14 Utility of combining CytoTRACE with Monocle 2 to identify rooted developmental hierarchies and lineage-specific genes without prior knowledge. (A-C)** Combined application of CytoTRACE and Monocle 2 to delineate complex branching processes during mouse bone marrow differentiation (Smart-seq2; Tabula Muris[42]) without prior knowledge of the root. (**A**) Unrooted tree predicted by Monocle 2 showing previously annotated bone marrow phenotypes[42]. (**B**) Pseudotime estimated by Monocle 2 after automatically determining the root with CytoTRACE. (**C**) Heat map of branch-specific genes identified by Branching Expression Analysis Modeling (BEAM)[34] using the CytoTRACE-rooted tree in panel **B**. Representative genes from each major hematopoietic lineage or group are shown. (**D-F**) Same as in **A-C** but applied to mouse intestine differentiation (Drop-seq)[43].

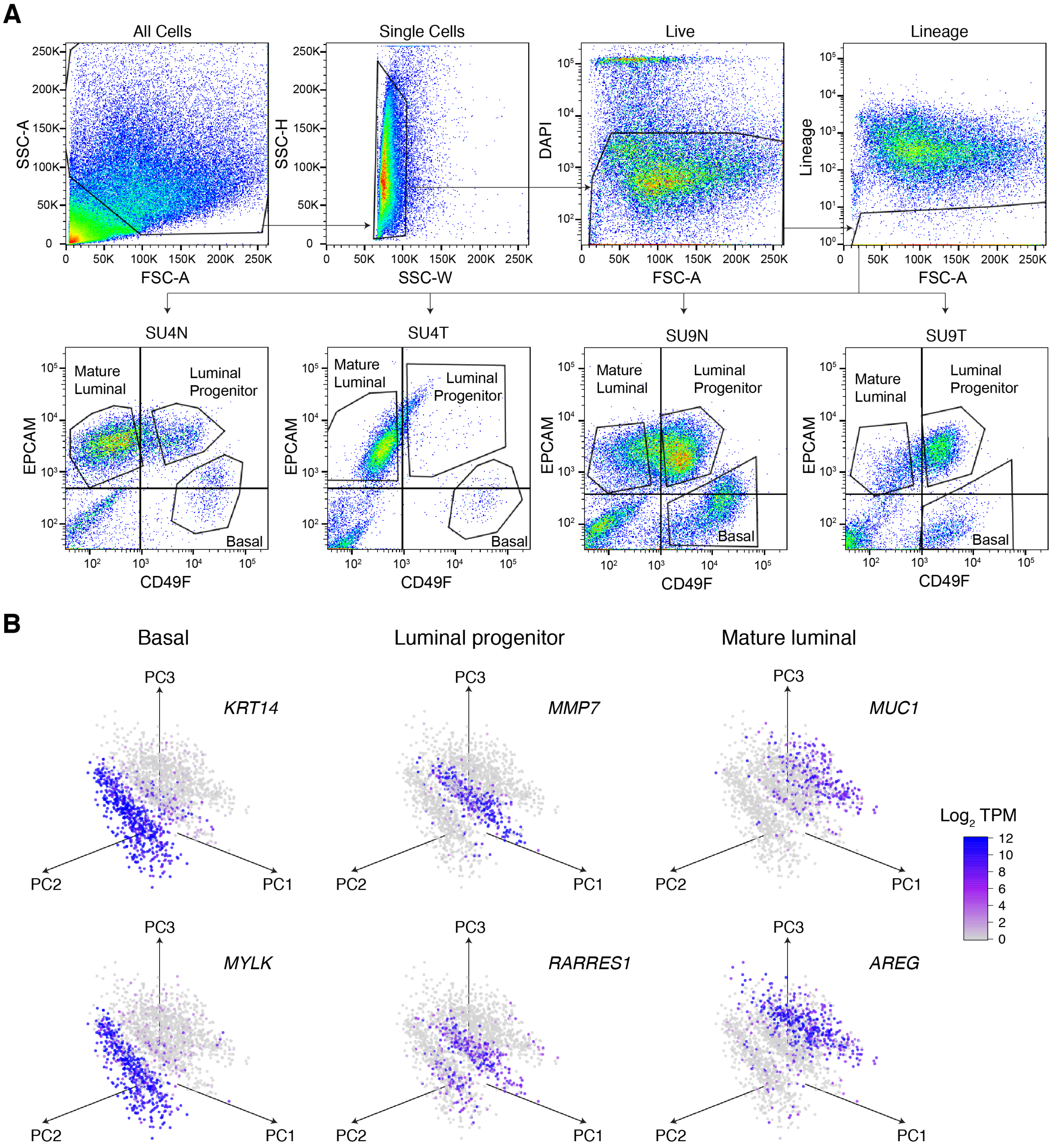


**Figure S15 Gating scheme and marker validation of epithelial subpopulations from human breast tumors and adjacent normal tissues.** (**A**) Gating scheme for the prospective isolation of normal and malignant human breast basal, luminal progenitor, and mature luminal cells. Representative profiles of paired normal and tumor samples from two representative patients, SU4 and SU9, are shown. (**B**) 3D plot of first three principal components showing gene expression patterns of marker genes for human breast basal (left; *KRT14* and *MYLK*), luminal progenitors (center; *MMP7* and *RARRES1*), and mature luminal cells (right; *MUC1* and *AREG*).

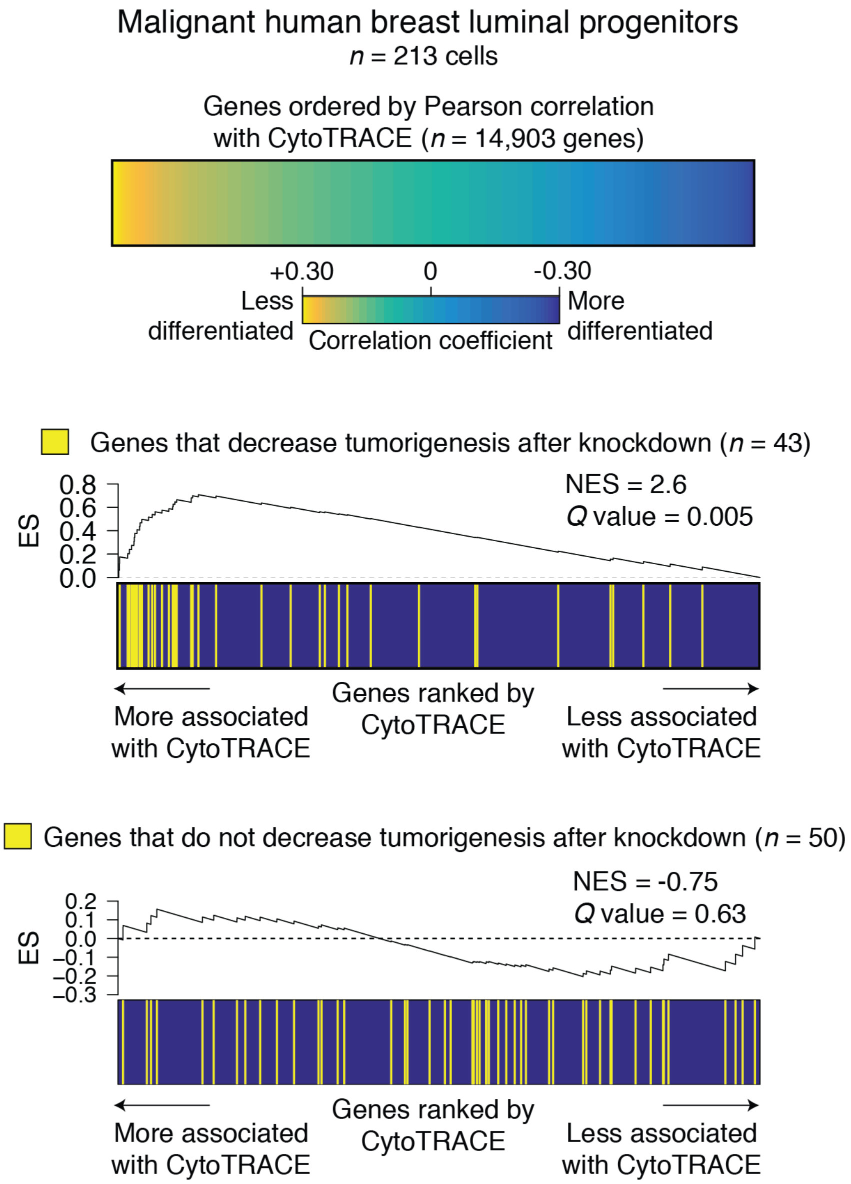


**Figure S16. Validation of CytoTRACE for prioritizing markers of breast tumorigenesis using results from an RNAi dropout viability screen.** Prediction of differentiation-associated genes in malignant luminal progenitors profiled by scRNA-seq. *Top:* Heat map showing malignant LP genes ordered by their Pearson correlation with CytoTRACE. *Center:* Pre-ranked gene set enrichment analysis[68] of 43 genes found to decrease human breast tumorigenesis in an RNAi dropout viability screen[57] in relation to LP genes ranked by CytoTRACE (same order as above). *Bottom*: Same as center panel, but with 50 genes found to have minimal effect on tumorigenesis after knockdown. NES, normalized enrichment score; ES, enrichment score.

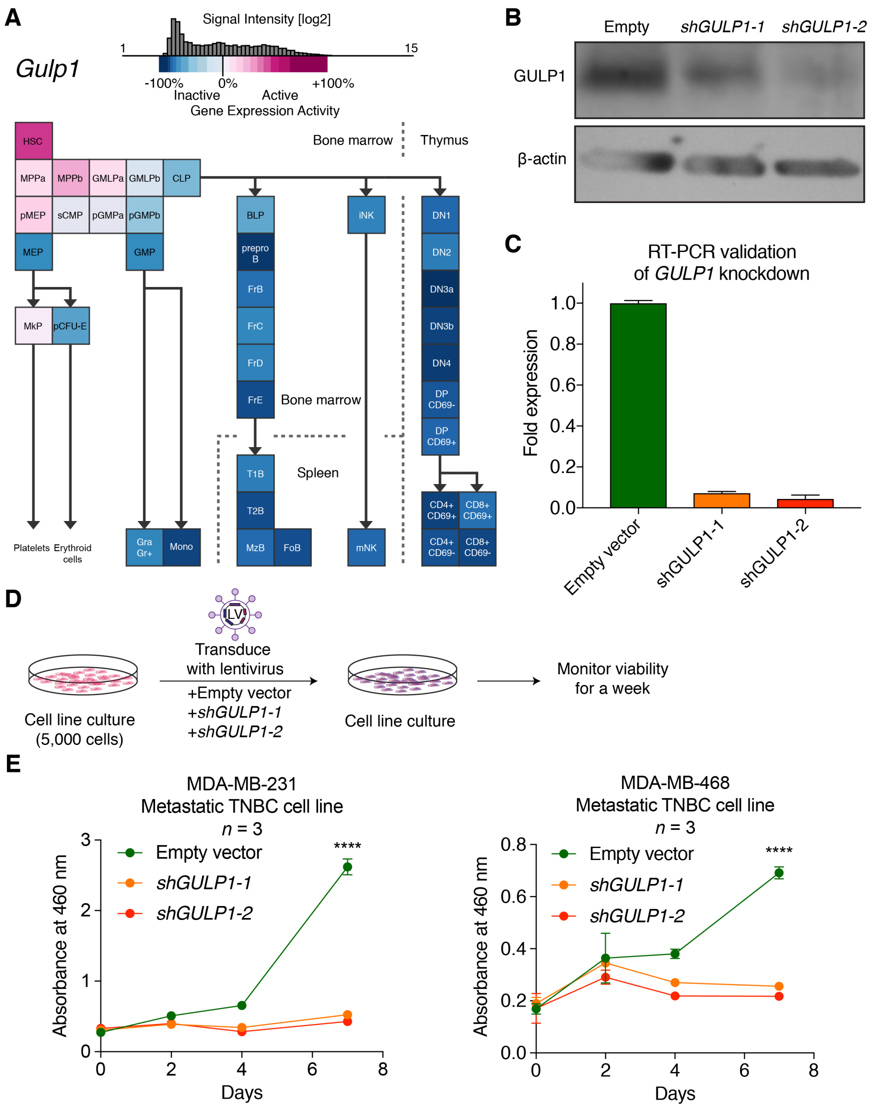


**Figure S17 Enrichment of *Gulp1* in hematopoietic stem cells and validation of *GULP1* knockdown.** (**A**) Heat map of *Gulp1* expression in the mouse hematopoietic stem cell tree, organized from the least differentiated cell, the hematopoietic stem cell (HSC), in the top-left to increasingly specialized progenitors and progeny towards the bottom. Each tile represents a cell type in hematopoiesis and the color indicates normalized gene expression of *Gulp1* (high expression = red; low expression = blue) as shown in the color legend at the top. Dotted lines separate cells from different compartments, e.g. thymus, spleen, or bone marrow. Figure panel was adapted from the 'Mouse Hematopoiesis Model' in Gene Expression Commons (http://gexc.riken.jp)[122]. (**B**) Western blot images of GULP1 protein expression after transduction of MDA-MB-231 cells with lentivirus containing an empty vector, *shGULP1-1, or shGULP1-2.* B-actin protein expression is shown below as housekeeping control. (**C**) Bar plot of fold expression change (y-axis) by real-time PCR (RT-PCR) in MDA-MB-231 cells after transduction with lentivirus containing an empty vector, *shGULP1-1*, or *shGULP1-2*. Fold expression change was normalized to empty vector control. (**D**) Schema for measuring cell viability after shRNA knockdown of *GULP1* in two metastatic TNBC cell lines, MDA-MB-231 and MDA-MB-468. (**E**) Viability of MDA-MB-231 (top; $n$ = 3) and MDA-MB-468 (bottom; $n$ = 3) after shRNA knockdown of *GULP1* as measured by change in absorbance at 460 nm ($y$-axis) over the course of a week ($x$-axis). Legend for color and conditions is indicated in the plot.
